# An ecological origin of the asymmetry of human color discrimination: evidence from psychophysics, MEG, and deep neural networks

**DOI:** 10.64898/2026.06.16.732654

**Authors:** Laysa Hedjar, Takuma Morimoto, Arash Akbarinia, Mandy V. Bartsch, Hendrik Strumpf, Jens-Max Hopf, Karl R. Gegenfurtner

## Abstract

Hue is a powerful cue for object discrimination, but human color discrimination is anisotropic: hue thresholds are lower than chroma thresholds for orangish colors, but nearly equal for purplish colors. Here, combining analyses of environmental color statistics, human psychophysics, magnetoencephalography (MEG), and deep neural networks, we provide converging evidence that this asymmetry reflects environmental color statistics, beyond the constraints imposed by cone-opponent architecture. Across fifteen image and reflectance databases, orangish hues accounted for 62.5% of chromatic samples, compared with 5.3% for purplish hues. We replicated the hue–chroma asymmetry psychophysically in 44 participants. MEG revealed a corresponding neural asymmetry, with superior decoding of orangish hue differences emerging around 250 ms after stimulus onset. Deep neural networks trained on naturalistic image datasets reproduced the human-like asymmetry without explicit color supervision. Critically, training on hue-inverted images reversed the asymmetry, producing greater hue sensitivity for purplish than for orangish colors. These results suggest that this long- known asymmetry in color discrimination is shaped by ecological chromatic structure.

## Introduction

Vision has evolved to solve the ecological problems organisms face. Perception reflects the regularities of the environment and the behavioral goals it serves, rather than passively recording the world (Gibson, 1979; Goodale & Milner, 1992; Milner & Goodale, 2006). Across species, sensory tunings align with ecologically relevant stimulus statistics, from orientation sensitivity in human vision to spectral sensitivities adapted to aquatic environments (Lythgoe & Partridge, 1989; Baden, 2024; Appelle, 1972; Furmanski & Engel, 2000; Girshick et al., 2011). Color vision offers a particularly revealing case of this principle. It arose in primates through an unusual evolutionary history in which trichromacy was reestablished through duplication and divergence of a long-wavelength opsin gene (Jacobs, 2009; Nathans et al., 1986; Surridge et al., 2003). This evolutionary detour, linked to the demands of frugivory (Osorio & Vorobyev, 1996; Mollon, 1989), resulted in a chromatic system characterized by asymmetric spectral sampling and recombination of cone signals into opponent mechanisms and higher-order cortical representations that support perception of hue and saturation (Krauskopf et al., 1982; Krauskopf et al., 1986; Gegenfurtner & Kiper, 1992; De Valois & De Valois, 1993; Kiper et al., 1997; Hansen & Gegenfurtner, 2006, 2013). Yet psychophysical measurements reveal systematic anisotropies in color discrimination that remain unexplained by cone-opponent architecture alone.

Hue is often more stable than chroma or luminance across changes in illumination and shading, making it a powerful cue for object identity (Gegenfurtner, 2025). Natural objects vary substantially in chroma and lightness but comparatively little in hue (*Figure 1*; Ennis et al., 2018; Klinker, Shafer, & Kanade, 1990; Ling, Vurro, & Hurlbert, 2008), leading to the long-standing proposal that hue should receive finer discriminative resolution than chroma (Judd, 1969). However, it has been shown that hue discrimination is markedly superior to chroma discrimination around colors appearing orange and blue, but not colors appearing purple and green (Krauskopf & Gegenfurtner, 1992; Giesel et al., 2009; Hansen et al., 2008; Danilova & Mollon, 2016; Hedjar et al., 2025a, 2025b). This anisotropy in color space has been repeatedly observed, yet its origin remains unresolved.

**Figure 1.**
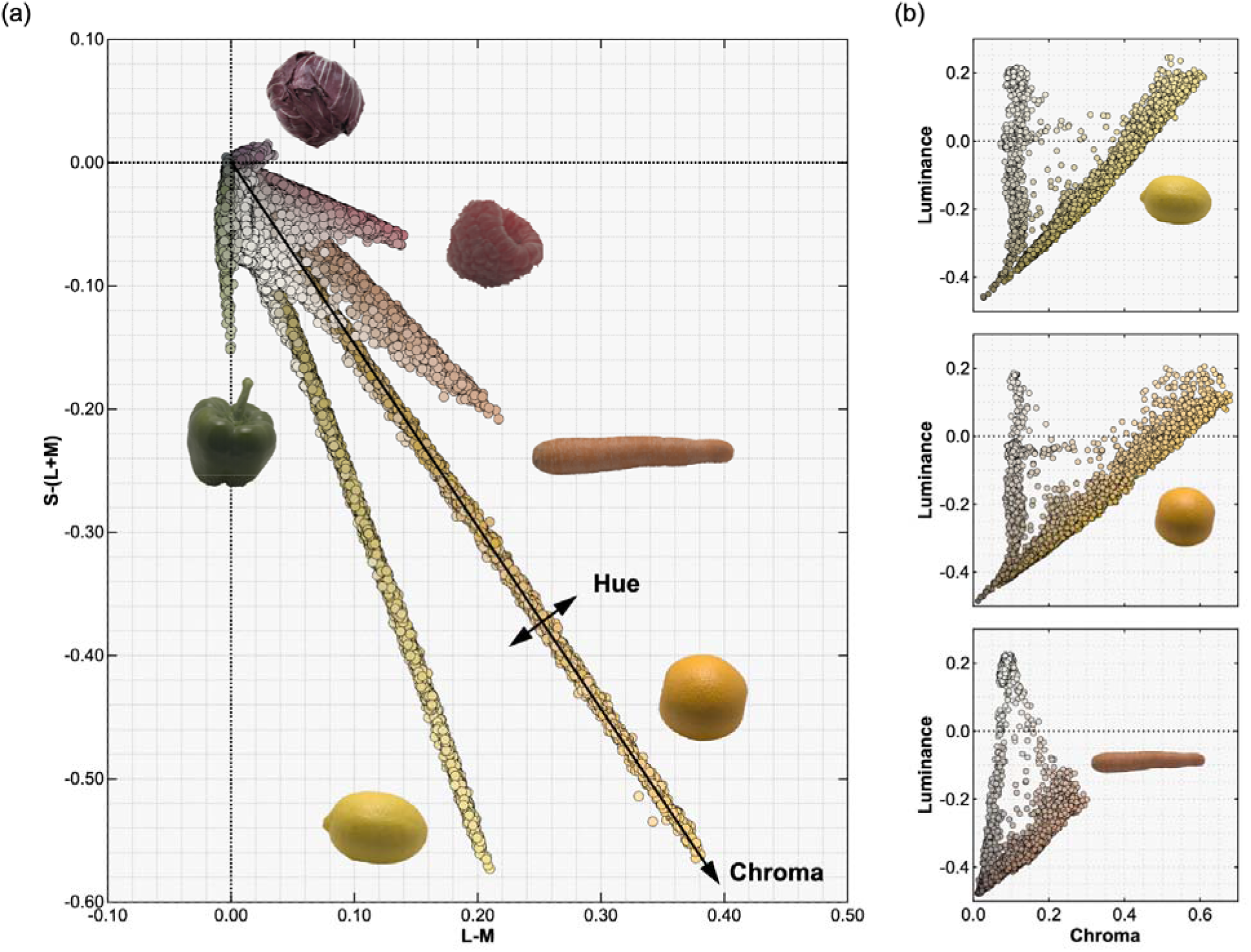
**Distribution of pixels from hyperspectral images of fruit and vegetable outer skins**. (a) Points are projected onto the isoluminant plane of DKL color space (Derrington et al., 1984). Arrow indicate the hue dimension (angular) and chroma dimension (radial). (b) DKL chroma is plotted against luminance for three objects, with data collapsed across hues. Both values are expressed relative to the monitor mid-grey. The high-luminance, low-chroma cluster corresponds to specular reflections. Data and object icons were taken from Ennis et al. (2018), except for the dark cabbage icon, which was obtained from PxHere available under a CC0 license. 5,000 pixels were randomly sampled from each object image to visualize the data density.

The idea that color vision is calibrated to the chromatic statistics of the environment has a long history. Natural color distributions are elongated along a bluish–yellowish axis close to the daylight locus (Webster & Mollon, 1997; Wachtler et al., 2001), and this structure has been linked to chromatic adaptation, the non-uniformity of color space, surface segmentation, the number of distinguishable colors, the perception of white, and aesthetic judgments (Webster & Mollon, 1997; McDermott & Webster, 2012; Fine et al., 2003; Marín-Franch & Foster, 2010; Bosten et al., 2015; Su et al., 2024; Wozniak et al., 2026). Danilova and Mollon (2010) found enhanced discrimination near the daylight locus and attributed it to natural illuminants, and color perception also tracks developmental exposure and visual diet (Skelton et al., 2023, 2024). This evidence has remained correlational, however, and has related perception to global color statistics such as the overall distribution of colors in natural scenes; how such global statistics translate into local, direction-specific sensitivities around particular colors has remained an open question. Addressing both issues requires a system in which the global color statistics of the environment can be manipulated while everything else is held constant, and that can, without explicit instruction, translate those statistics into its own color sensitivity.

An ecological account predicts that this asymmetry of color discrimination should reflect the structure of natural color distributions. Directly testing this hypothesis in humans has been hampered by the difficulty of manipulating visual experience over the prolonged developmental timescales on which such sensitivities are likely to emerge. Here, we overcome this challenge by combining analyses of environmental color statistics, human psychophysics, cortical dynamics, and task-optimized deep neural networks. These networks provide an experimentally tractable system in which the color statistics available during learning can be systematically manipulated. By training on natural images, the networks acquire relevant regularities without direct supervision for color discrimination. This approach allows us to test whether natural color statistics shape a long-standing asymmetry in human color discrimination, whether this asymmetry is selectively expressed in task-relevant cortical activity, and whether it can be reversed by altering the color statistics experienced during learning.

## Results

### Distribution of colors in the world is biased towards orange and blue

We first characterize the non-uniformity of hue distributions in the physical environment by analyzing multiple datasets compiled for diverse purposes including object recognition, segmentation, and concept curation. *Figure 2* plots hue histograms across natural surface reflectances, uncalibrated RGB images, calibrated RGB images, and hyperspectral images in the physiologically-based color space DKL (Derrington et al., 1984). Despite substantial differences in acquisition methods and image content, a remarkably consistent pattern emerged: the chromatic world is highly asymmetric. The orange quadrant contained the largest share of chromatic samples, followed by a secondary peak on the blue side, while the purple quadrant was sparsely populated. These orange hues also encompass their darker counterpart, brown—a hallmark of natural substrates such as soil and other earthy materials. Contrary to intuition, vegetation, including grass and leaves, fell largely along the y-axis and in the orange quadrant rather than in the nominally green quadrant. Formally, treating each dataset as one observation, the mean fraction of samples in the orange quadrant across datasets was 62.5%, whereas the purple quadrant accounted for only 5.3%; this difference was statistically significant (two-tailed paired *t*-test across datasets: *t*(14)=14.2, *p*=1.05×10^-9^; Cohen’s *d*=3.79). Because the number of images differs substantially across datasets, we repeated the test weighting each dataset by its image count, with degrees of freedom based on Kish’s effective sample size (effective n = 2.5, reflecting the dominance of Places365 and ImageNet). This confirmed the same trend (orange 59.1%, purple 4.4%; *t*(1.5)=25.1, *p*=5.68×10 ³, Cohen’s *d*=15.8).

**Figure 2.**
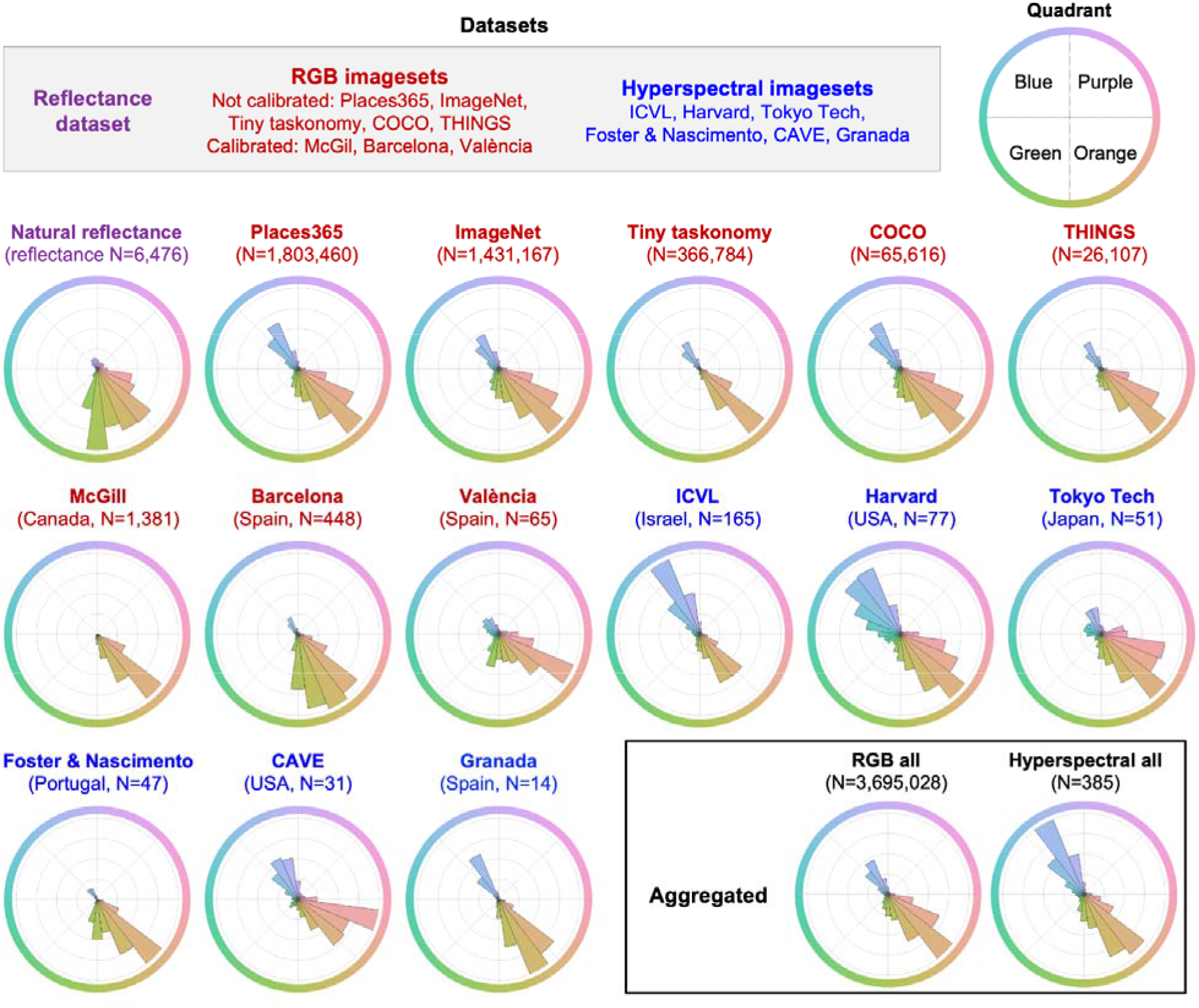
DKL hue histograms of pixels across images or reflectances in each database. N indicates the number of reflectances for the natural reflectance dataset and the number of evaluated images for the other datasets. In addition to the histograms for individual datasets, two additional histograms show the distributions aggregated across all RGB image databases and across all hyperspectral image databases. The country of data collection is noted where available. In the top right, we assign color terms to the four DKL quadrants based on the approximate appearance of their colors.

This predominance of orange hues is consistent with the observation that many biologically relevant objects (including skin, soil, fruit, and vegetation) occupy this region of color space, whereas blues are often associated with spatially extended backgrounds such as sky and water (Rosenthal et al., 2018). Thus, the ecological color environment is not chromatically balanced, but strongly biased toward the orange–blue axis, with a pronounced underrepresentation of purple hues.

### Psychophysical evidence of hue superiority for orange but not purple colors

The strong contrast between the prevalence of orange relative to purple in the environment led us to ask whether human discrimination exhibits a similar asymmetry. If perceptual sensitivity reflects natural color distributions, we would expect enhanced discrimination for hue variations in the orange region of color space, where such variations are common, compared to the more sparsely populated purple region.

To test this prediction, we measured and compared discrimination thresholds along hue and chroma directions in DKL color space for orange and purple colors. This local comparison asks whether discrimination is preferentially enhanced along the hue direction, rather than reflecting overall sensitivity to chromatic variation within each color region. In a four-alternative forced- choice task, participants identified the “odd one out” among four colored discs, where one disc differed from the others either in hue or in chroma. Thresholds were estimated separately for orange and purple reference colors and converted to sensitivities by taking their reciprocal.

The results closely replicated previous findings (Danilova & Mollon, 2016; Giesel et al., 2009; Hedjar et al., 2025a, 2025b; Krauskopf & Gegenfurtner, 1992). For orange colors, participants were substantially more sensitive to hue than to chroma differences, whereas for purple colors hue and chroma sensitivities were much more similar (*Figure 3*). Consequently, hue–chroma sensitivity ratios were significantly larger for orange than for purple colors (mean log_10_ ratios: orange = 0.251, purple = 0.061; two-tailed *t*(43)=15.19, *p*=6.77×10⁻¹, 95% *CI* difference between ratios = [0.164, 0.215], Cohen’s *d*=2.29).

**Figure 3.**
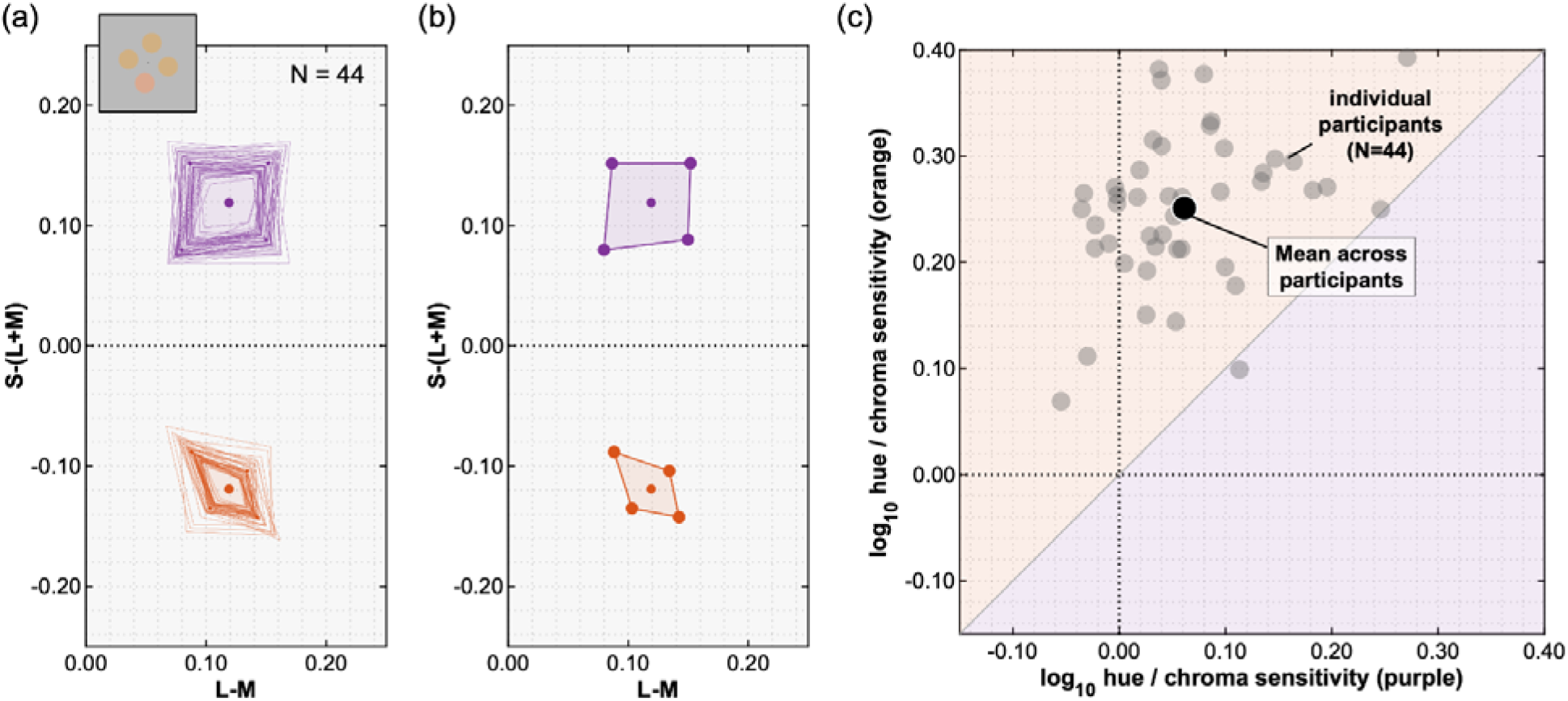
Hue and chroma discrimination thresholds and sensitivities for purple and orange colors. (a) Thresholds plotted on the orange and purple quadrants of the isoluminant plane of DKL color space for 44 individual participants. Reference colors are plotted in orange and purple and thin lines connect the thresholds for an individual. The top-left inset shows the stimulus arrangement in an example trial. (b) Thresholds averaged across all 44 participants. (c) Hue-to-chroma sensitivity (1/threshold) ratios for each participant plotted on a logarithmic scale, with purple ratios on the x-axis and orange on the y- axis. We averaged thresholds across the two polarities per color direction (e.g., increasing and decreasing chroma) before computing sensitivity. The dotted vertical and horizontal lines show equal hue and chroma sensitivities.

The magnitude of this effect was striking: only the orange region showed a pronounced preference for hue over chroma discrimination. This asymmetry closely mirrors the ecological imbalance shown in *Figure 2*, where orange hues dominate natural color distributions while purple hues are rare. Thus, the structure of human color discrimination appears to reflect the large-scale chromatic regularities of the natural environment. This correspondence is, of course, correlational; a more direct test of whether environmental statistics can produce the asymmetry is provided by the network experiments below. Nevertheless, since a primary function of color vision is to facilitate object identification and discrimination, enhanced sensitivity to differences in the orange region likely confers an adaptive advantage in recognizing biologically relevant objects.

### Physiological responses to color discrimination consistent with psychophysical sensitivities

We next asked whether the psychophysical hue–chroma asymmetry was reflected in cortical activity. Participants performed the same four-alternative odd-one-out color discrimination task during magnetoencephalography (MEG) recording. To relate neural responses directly to behavior, we used each participant’s psychophysical thresholds to select individualized chromatic shift magnitudes. On each trial, the odd disc was either identical to the orange or purple reference or shifted from it in hue or chroma by a small, medium, or large amount.

Within each participant, these shift magnitudes were physically matched across color directions: for each magnitude, colors were equidistant from their reference in DKL space and produced matched changes along the cardinal chromatic dimensions. This design allowed us to compare neural discriminability across color directions at equivalent physical DKL distances, even though their perceptual discriminability differed. Because orange hue thresholds were markedly lower than the other thresholds, orange hue shifts were generally suprathreshold, while purple hue, purple chroma, and orange chroma shifts sampled a broader range of the psychometric function (see *Supplementary Information (SI)*, *Figures S1* and *S2*). This difference in perceptual discriminability was central to our prediction: if MEG activity reflects psychophysical sensitivity, we would expect decoding to reveal a stronger hue advantage for orange than for purple colors.

#### Decoding hue and chroma differences from MEG responses

We analyzed whether the psychophysical asymmetry between hue and chroma discrimination was reflected in cortical activity. Using single-trial MEG responses, we trained classifiers to distinguish stimuli containing a hue- or chroma-shifted odd disc from stimuli in which all discs were identical. Separate classifiers were trained for small, medium, and large color shifts relative to the orange and purple reference colors.

*Figure 4a-d* plots the time course of decoding accuracy for hue and chroma differences around orange and purple reference colors. A clear asymmetry emerged: for orange colors, hue differences were decoded substantially better than chroma differences across all shift magnitudes, whereas for purple colors the difference was much smaller. Decoding accuracy was therefore more strongly related to psychometric performance than to chromatic distance (see also *Figure S3*). This neural asymmetry closely mirrored the psychophysical pattern shown in *Figure 3*.

**Figure 4.**
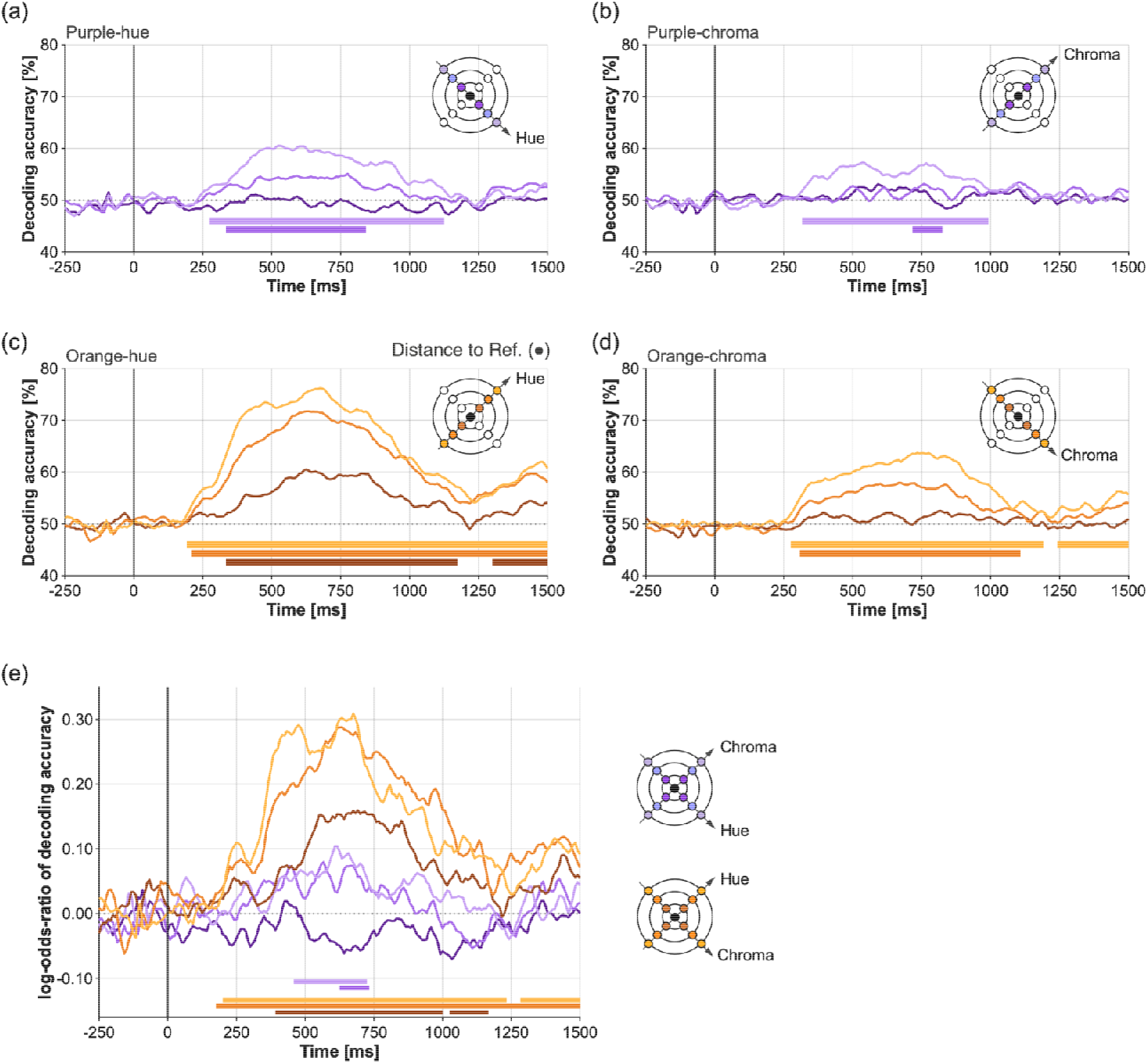
Multivariate pattern analysis of the odd disc color shifts for hue and chroma. (a-b) Average decoding accuracy (% correct classification) for purple colors with small (plotted as a dark purple line), medium (purple line), and large (light purple line) odd disc shifts along (a) hue and (b) chroma directions. The horizontal bars indicate the time range of significant decoding (Monte Carlo significance probability p<.05) obtained with cluster-based permutation testing. (c-d) Average decoding accuracy for orange colors with small (plotted as a brown line), medium (orange line), and large (light orange line) odd disc shifts along (c) hue and (d) chroma directions. (e) Hue-to-chroma log-odds-ratio of decoding accuracy for the three shift conditions of orange and purple colors. Time zero denotes stimulus onset in panels (a-e). Stimuli were displayed for 500 ms.

Decoding accuracy increased with chromatic distance from the reference, and larger shifts produced earlier and more reliable decoding. Importantly, the hue advantage for orange colors emerged consistently after approximately 250 ms post-stimulus onset and was evident across all shift magnitudes (*Figure 4e*). Thus, the cortical representation of color differences exhibited the same selective enhancement of hue discrimination in the orange region of color space that was observed behaviorally. Source localization (see *SI*, *Figure S5*) suggests that the response evolved from early visual cortex toward a distributed ventral occipital network encompassing regions associated with color processing and visual selection.

To compare neural and behavioral asymmetries directly, we plotted psychophysical and MEG- derived hue–chroma ratios in a common coordinate system (*Figure 5a*). For the largest shifts, later MEG responses clustered near the psychophysical group mean, indicating that the neural representation increasingly approximated the behavioral pattern over time. Further analyses confirmed that the MEG-derived ratios progressively approached the psychophysical group mean as shift magnitude increased across small, medium, and large shifts (*Figure S4a-c*).

**Figure 5.**
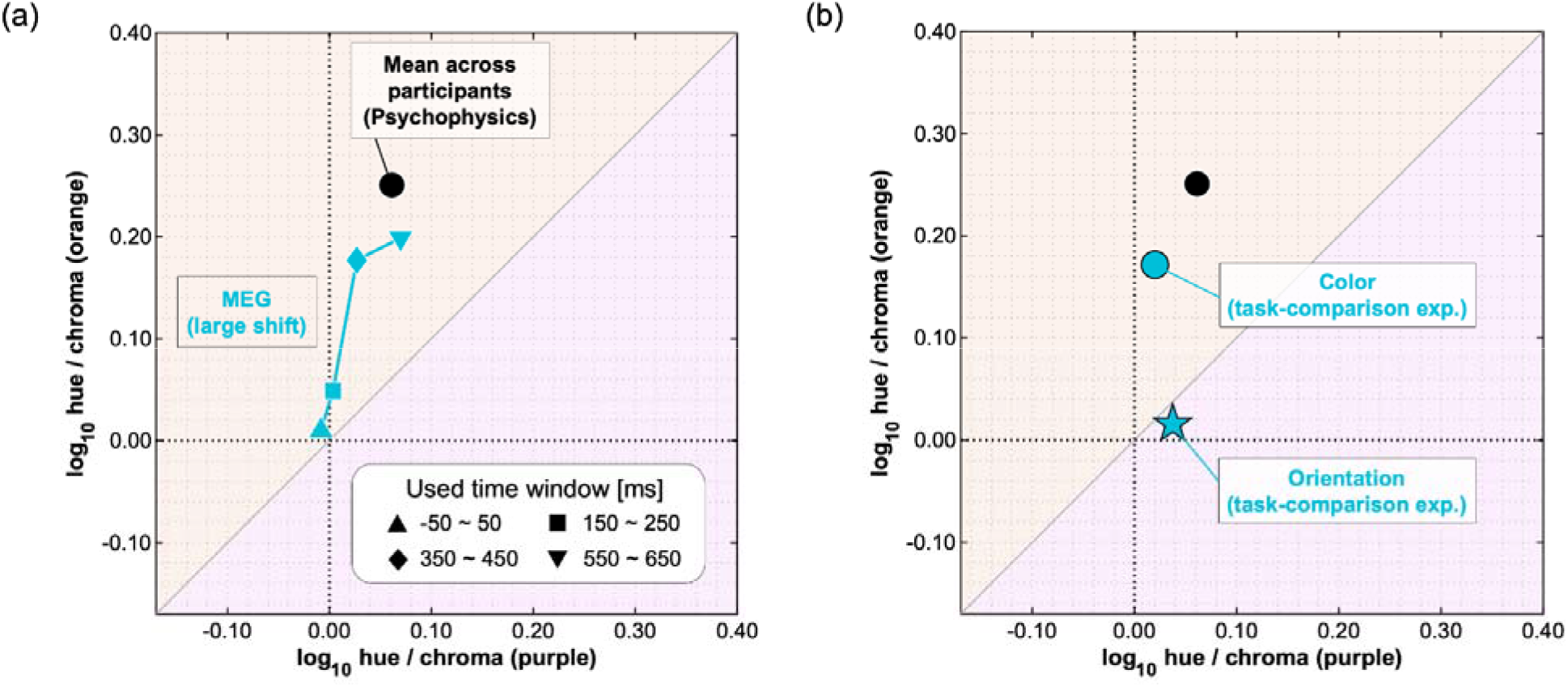
Relationship between psychophysical hue superiority and MEG-derived hue–chroma decoding asymmetries for purple and orange colors. (a) MEG-derived hue-to-chroma log-odds-ratios are plotted against the psychophysical hue-to-chroma ratios, with purple on the x-axis and orange on the y-axis. The black circle shows the psychophysical group mean, replotted from Figure 3. Cyan symbol show MEG group means for the large odd disc color shift across different post-stimulus time windows, with symbol shape denoting the time window. Later MEG responses moved closer to the psychophysical group mean, indicating that the neural decoding asymmetry became more similar to the behavioral pattern. (b) Hue-to-chroma log-odds-ratios from the task-comparison MEG experiments, averaged between 350 and 650 ms post-stimulus onset for the large color shifts. The black circle shows the psychophysical group mean. The cyan circle shows the group mean for the color discrimination task, and the cyan star shows the group mean for the orientation discrimination task. The color discrimination tas showed a hue–chroma asymmetry closer to the psychophysical pattern than the orientation discrimination task.

#### Task-dependent MEG decoding of chromatic differences

Having established that MEG decoding reproduced the psychophysical asymmetry between orange and purple colors, we next asked whether this neural asymmetry was expressed whenever chromatic information was present or whether it depended on the behavioral relevance of color. Participants viewed the same stimuli in two separate sessions. In one session they performed the same color discrimination task as before. In the other, they performed an orientation discrimination task and reported whether the stimulus was tilted clockwise or counterclockwise. Thus, the visual input was identical while task demands differed.

Using the same decoding procedure as in the main experiment, we asked whether responses to hue- or chroma-shifted stimuli could be decoded from responses to non-shifted stimuli. *Figure 5b* summarizes the hue-to-chroma decoding asymmetry for the two tasks. As in the main MEG experiment, the color discrimination task showed ratios similar to the psychophysical data, with greater hue sensitivity for orange over purple colors (two-tailed one-sample *t*-test on signed distances to unity line: mean log-odds-ratio distance = 0.107 on orange side, 95% *CI* [0.072, 0.142], *t*(26) = 6.30, *p* = 1.14×10^-6^, Cohen’s *d* = 1.21; see *SI*, *Figure S4d*, for individual participant data). In contrast, notably, decoding in the orientation task did not differ significantly from chance in any condition. No reliable asymmetry was observed; neither the orange nor the purple hue-to-chroma ratios differed significantly from zero (two-tailed one-sample *t*-tests against 0; purple: mean log-ratio = 0.037, 95% *CI* [-0.010, 0.085], *t*(18) = 1.64, *p* = 0.118; orange: mean log-ratio = 0.016, 95% *CI* [-0.013, 0.045], *t*(18) = 1.14, *p* = 0.271). Full decoding time courses are shown in *Figures S6 and S7*.

Thus, chromatic differences that were clearly expressed in the MEG signal during active color discrimination were not reliably expressed when participants judged orientation instead. Because the visual stimuli were identical across tasks, this finding suggests that task relevance, rather than stimulus discriminability alone, determined whether the ecological hue–chroma asymmetry was reflected in cortical activity.

### Neural networks trained on natural images spontaneously elicit discrimination thresholds similar to humans

If the hue–chroma asymmetry reflects environmental color statistics, then visual systems trained on natural images should acquire a similar pattern even without explicit supervision about color discrimination. To test this idea, we estimated chromatic discrimination thresholds in deep neural networks trained for object recognition, scene categorization, object detection, and pose estimation using image sets that exhibited a hue bias (*Figure 2*). Thresholds were measured using a linear-probe odd-one-out paradigm closely matched to the human psychophysical task (see *Methods*).

Networks trained on chromatically biased image datasets consistently reproduced the human pattern of color discrimination. ResNet50 (He et al., 2016) trained on the ImageNet dataset (Deng et al., 2009) for object recognition exhibited a clear hue superiority in the orange region of color space, whereas hue and chroma sensitivities were much more similar in the purple region (*Figure 6, upper left*). A nearly identical pattern emerged in networks trained for scene categorization (lower left) on Places365 (Zhou et al., 2017), and similar results were observed across architectures, datasets, and visual tasks (*Figure 7; Figure S8*). Layer-wise analyses showed that alignment with human sensitivity emerged in early layers for most visual tasks (*Figures S9 and S10*).

**Figure 6.**
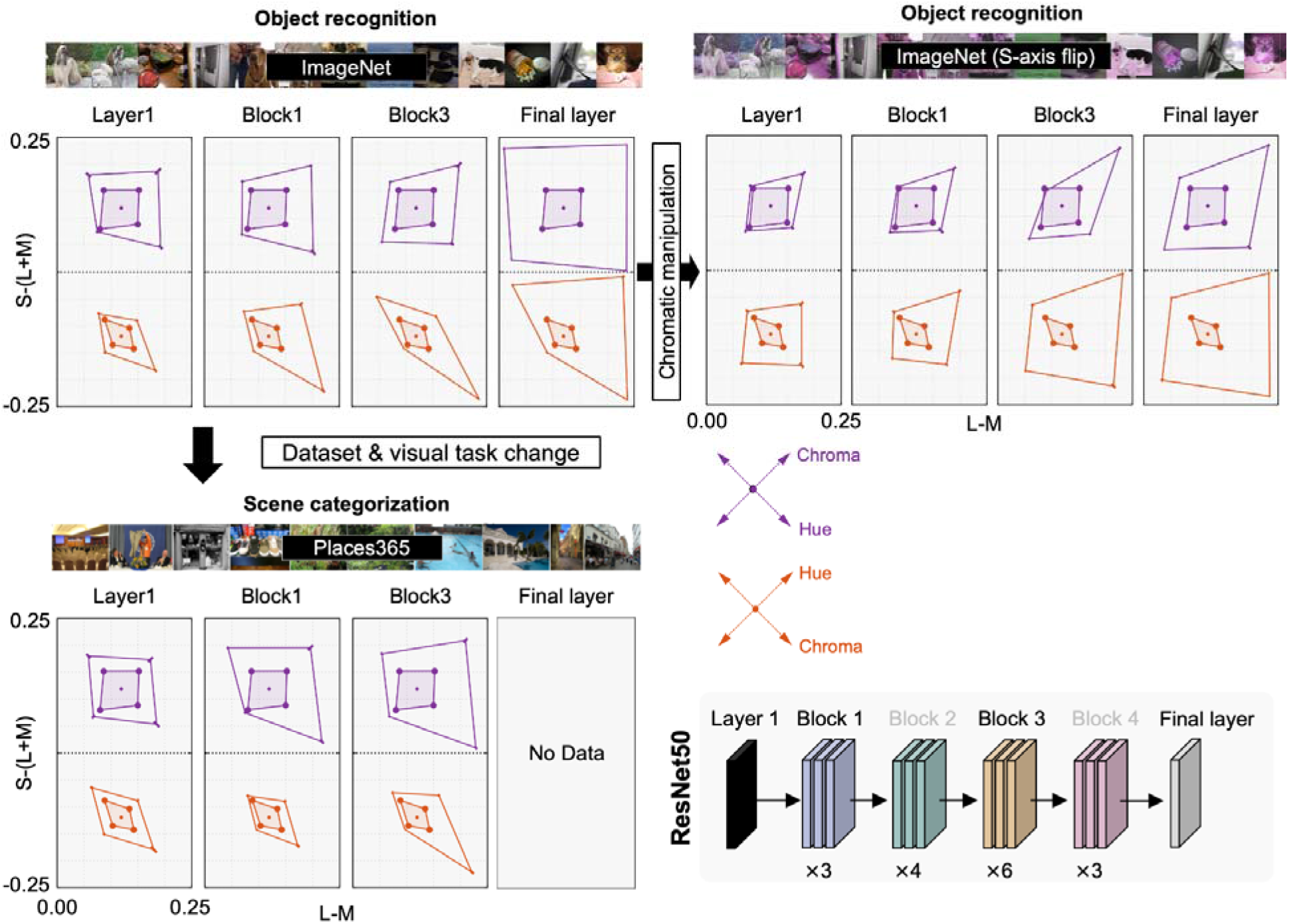
Discrimination thresholds in the orange and purple quadrants across processing stages of ResNet models. Threshold contours are shown in DKL color space for three networks: (upper left) ResNet50 trained on ImageNet for object recognition, (lower left) ResNet50 trained on Places365 for scene categorization, and (upper right) ResNet50 trained on an S-axis-flipped version of ImageNet for object recognition. Each column shows thresholds estimated from a different processing stage, from the initial layer through successive residual blocks and the final classification layer where available as shown in the lower right. Purple and orange contours indicate thresholds around the purple and orange reference colors, respectively; averaged human thresholds are shown by the inner contours and networ thresholds by the outer contours. For S-axis-flipped ImageNet, out-of-gamut colors were brought into gamut by shortening their DKL vectors, primarily reducing chroma and luminance while preserving hue. Each threshold represents an average across 10 iterations with different random seeds; error bars indicate ± 1 standard error.

**Figure 7.**
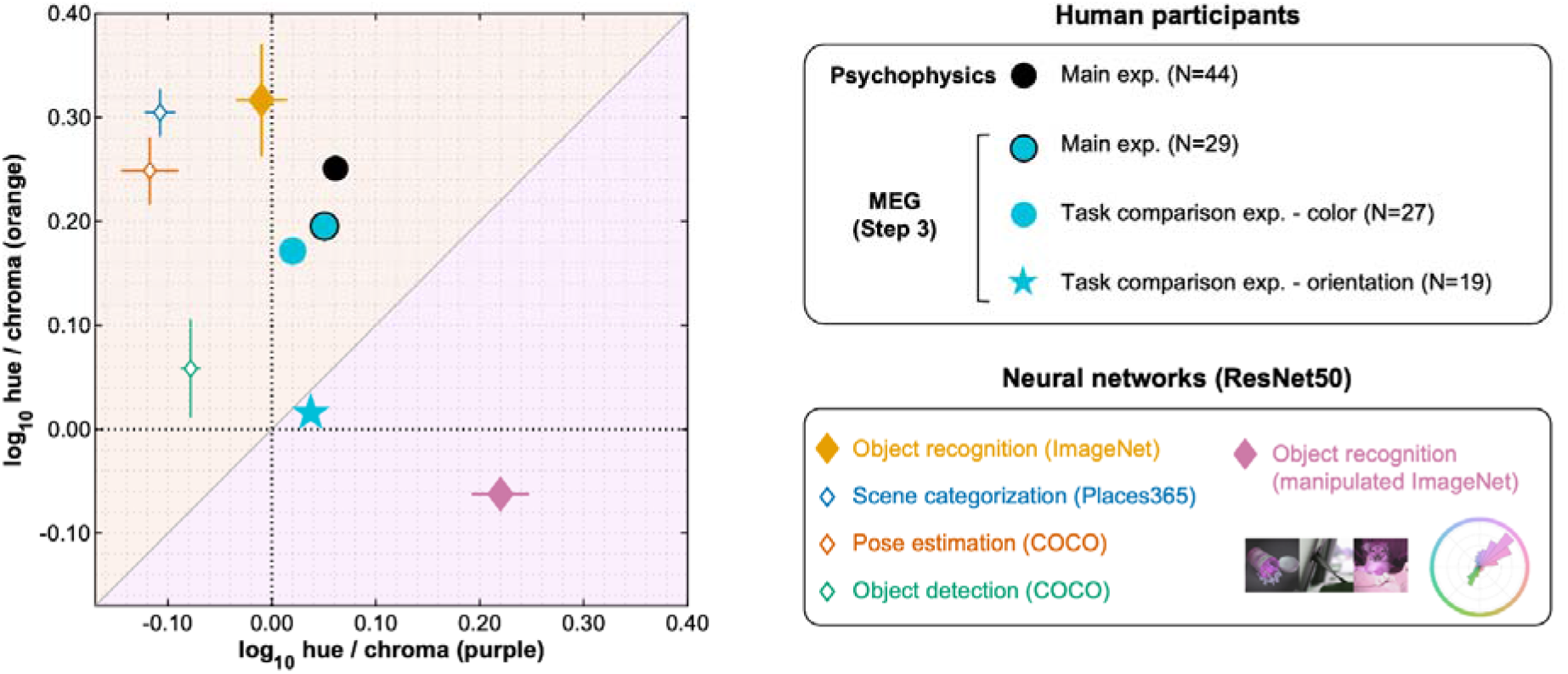
Comparison of the hue-superiority metric across different domains (psychophysics, MEG, and neural networks), shown for purple colors (x-axis) and orange colors (y-axis). For the psychophysical data and network outputs, hue-to-chroma sensitivity was computed as the inverse of threshold (1/threshold) and log-transformed. For MEG, we plotted hue-to-chroma decoding accurac odds-ratios derived from MEG signals at the large shift averaged over 350–650 ms after stimulus onset, also on a logarithmic scale. The black circle indicates the group mean of the psychophysical data, replotted from Figure 3. The cyan circle with a black outline indicates the group mean of the MEG data from the main color discrimination experiment. The cyan circle without an outline and cyan star represent the group means of the MEG data from the task-comparison experiment (color and orientation discrimination, respectively; replotted from Figure 5). Neural network results are shown for ResNet50 (diamonds). Colors indicate the training objective: object recognition, scene categorization, pose estimation, and object detection. Network data points were obtained by averaging across 10 iterations and then across processing depths. Error bars for the networks indicate ±1 S.E. across processing depths.

The strongest test of the ecological hypothesis came from manipulating the chromatic statistics of the training environment. We trained ResNet50 from scratch on a modified version of ImageNet in which the sign of the S–(L+M) axis was inverted, making purplish hues common and orangish hues rare. This manipulation reversed the discrimination asymmetry (*Figure 6, upper right*): hue superiority disappeared in the orange region and instead emerged around purple colors. Because network architecture, optimization procedure, and task objective remained unchanged, the reversal can only be attributed to the altered chromatic statistics of the training images. Thus, the asymmetry is not an intrinsic property of the network architecture, but a learned consequence of the visual environment.

To quantify hue superiority, we computed hue-to-chroma sensitivity ratios for all networks and compared them with the psychophysical and MEG data (Figure 7). Most networks trained on images with natural image statistics occupied the same region of the plot as human observers, exhibiting stronger hue superiority for orange than purple colors. The only exception was the object recognition network trained on the purple-dominant ImageNet, which shifted toward the opposite quadrant and showed enhanced hue sensitivity for purple colors. Within the network framework, this reversal provides causal evidence that the asymmetry is not determined by network architecture or task demands alone, but is shaped by the chromatic statistics of the training environment.

Remarkably, none of the networks received explicit supervision about color discrimination or hue–chroma relationships. Optimized solely for visual tasks such as object recognition, scene categorization, object detection, and pose estimation, they nevertheless developed color discrimination asymmetries that closely mirror those of human observers. Furthermore, because the networks were frozen and the readout was trained on colors sampled uniformly from RGB space, the readout could not introduce a chromatic bias; the asymmetries reflect information already present in the pretrained representations (see Methods).

## Discussion

One of the key questions in neuroscience is how the statistical structure of the external world becomes embedded in perception and its neural implementation. Human color discrimination provides a fruitful test case for this question: sensitivity is highly non-uniform across color space, and recent work has continued to characterize this anisotropy in increasing detail (Hong et al., 2025; Koenderink et al., 2026). Yet, what this asymmetry ultimately stems from has remained unanswered. Here, by combining access to over a million real-world color images with powerful learning systems capable of efficiently extracting chromatic regularities from those large-scale visual data, we show how natural scene statistics can shape a fundamental anisotropy of the color vision system. In humans, hue–chroma discrimination sensitivity ratios vary systematically across color space: around orange colors they are markedly larger than around purple colors (Danilova & Mollon, 2016; Giesel et al., 2009; Hedjar et al., 2025a, 2025b; Krauskopf & Gegenfurtner, 1992). Using MEG, we identified a corresponding pattern in the decodable neural responses emerging around 250 ms after stimulus onset in brain regions implicated in color selection. This perceptual and neural anisotropy mirrors the chromatic structure of the natural environment: across diverse image and reflectance databases, orange hues occur far more frequently than purple, consistent with the notion that the visual system, constrained by limited neural resources, allocates finer discriminative resolution to the signals it encounters more often. To directly test causality, we used deep neural networks (DNNs) trained on a variety of everyday tasks on chromatically biased natural images and measured their resulting hue– chroma sensitivities. Nearly all networks reproduced the human-like asymmetry despite no explicit supervision on color. Critically, when we trained an object recognition network from scratch on an image set with the opposite hue bias (purple dominating), the asymmetry reversed: hue sensitivity exceeded chroma sensitivity more for purple than for orange colors.

A growing body of work suggests that color vision is shaped by the statistics of natural scenes (see Introduction). Our results extend this literature in three respects: the environmental regularity we identify is a pronounced orange–purple imbalance in hue prevalence, consistent across fifteen datasets, rather than the familiar bluish–yellowish elongation; this imbalance is linked to a specific, long-known asymmetry in discrimination that is also evident in task- dependent cortical activity; and, most importantly, the network experiments go beyond correlation by showing that altering hue statistics during learning, with architecture, objective, and training procedure held constant, is sufficient to reverse the asymmetry.

The hue imbalance likely reflects variation in both natural surfaces and natural illumination. Daylight varies from bluish skylight to yellowish-orange sunlight along a locus that runs through the blue and orange quadrants of the DKL plane, so illumination adds chromatic variation mainly along the orange–blue axis. For conventional image datasets collected from the internet, such as ImageNet, however, individual images have likely undergone white balancing and other camera processing, making the contribution of illumination difficult to estimate directly. The reflectance datasets, comprising a subset of the hyperspectral images and rendered under a single D65 illuminant, nevertheless showed the same predominance of orange. Thus, illuminant variation would reinforce rather than counteract the pattern, concentrating chromatic variation in the orange and blue quadrants while leaving purple scarce, consistent with proposals that discrimination is enhanced near the daylight locus (Danilova & Mollon, 2010) and that vision incorporates a prior for natural daylights (Bosten et al., 2015; Su et al., 2024).

Our MEG results extend this ecological account by revealing a corresponding neural asymmetry. Hue differences in the orange region of color space were decoded more accurately than chroma differences, closely mirroring the psychophysical results. Previous MEG and EEG studies have demonstrated reliable decoding of stimulus color under passive viewing, weakly task-constrained viewing, and active feature-report paradigms (Rosenthal et al., 2021; Hajonides et al., 2021; Hermann et al., 2022; Chauhan et al., 2023). Goddard and Mullen (2025) also found that perceptual color differences can explain nonuniformities in neural color representations beyond those captured by a cone-contrast model (see also Rosenthal et al., 2021; Chauhan et al., 2023). Much of this literature, however, has primarily focused on decoding the color of individual stimuli or remembered items. An important exception is Macyczko et al. (2026), who showed that asymmetries in cortical responses to hue differences mirror perceptual asymmetries, despite equivalent hue separations in cone-opponent color space. During VEP recording, however, participants passively viewed stimuli with suprathreshold hue differences. In contrast, the present experiment required participants to actively discriminate subtle, within-display chromatic differences during recording, including differences near psychophysical threshold. We also used individually tailored chromatic shifts matched in physical DKL distance and performed participant-specific decoding. The demands of fine discrimination may also be relevant to the timing of the neural effect: the decoding asymmetry emerged relatively late, beginning around 250 ms after stimulus onset. This timing is consistent with previous findings that coarse color information becomes available earlier, whereas fine chromatic discrimination requires additional recurrent processing in order for representations to reach sufficient resolution (Bartsch et al., 2017; Hermann et al., 2022).

Critically, the asymmetry disappeared when the same stimuli were viewed during an orientation discrimination task. Thus, chromatic information was not automatically expressed whenever it was present in the stimulus, but rather depended on the behavioral relevance of color. This does not imply that early sensory representations contain no chromatic asymmetry, since weak feedforward signals may become measurable only when amplified by task demands (Bartsch et al., 2017). Nevertheless, the combination of late decoding onset, involvement of color- responsive mid-level VVC regions (see *SI*, *Figure S5*), and the absence of reliable decoding during orientation judgments suggests that the observed hue–chroma asymmetry was expressed through task-dependent cortical dynamics engaged during active color discrimination. More broadly, the findings are consistent with evidence that color representations in higher visual cortex depend on behavioral goals and task context (Koida & Komatsu, 2007; Kuriki et al., 2025), as well as with a growing literature emphasizing task- dependent neural coding across sensory systems (Nau et al., 2024).

Feature-based attention provides a plausible mechanism for this task dependence (Bartsch et al., 2015; Jehee et al., 2011; Müller et al., 2006; Huk & Heeger, 2000; Schulz et al., 2024), but is unlikely to explain the specific structure of the asymmetry on its own. Attention to color can sharpen population tuning through recurrent processing in the visual hierarchy (Bartsch et al., 2017; Schulz et al., 2024), making fine chromatic differences more separable when color is task-relevant. However, generic attentional sharpening does not explain why this improvement should preferentially favor hue over chroma in the orange region of color space. Since all stimulus conditions appeared equally often, there is no obvious reason why attention should selectively benefit one chromatic direction over another. A more plausible account is that feature-based attention operates on an underlying representational geometry shaped by long- term exposure to environmental color statistics. Under this view, ecological structure determines which chromatic distinctions are represented with higher resolution, whereas active discrimination determines when these distinctions are selectively amplified and become decodable.

The deep neural networks reproduced the qualitative structure of the human hue–chroma asymmetry. However, several quantitative differences remain informative. In particular, DNN hue–chroma sensitivity ratios occupied a narrower range than those observed in human observers, and hue thresholds were often less symmetrical than the corresponding psychophysical thresholds. These discrepancies likely reflect differences between natural human visual experience and the statistics available during network training, as well as differences in architecture and learning objectives. We therefore view neural networks not as quantitative replicas of human perception, but as controllable models that allow causal tests of how environmental statistics shape color discrimination.

Consistent with this interpretation, extending the analysis across a broader range of visual objectives showed that human–network alignment depends more on environmental color statistics than on any particular task (see *SI*). Across a single ResNet50 architecture trained on 24 different Taskonomy objectives (Zamir et al., 2018), almost every network reproduced the human-like orange hue superiority regardless of its task, with the closest match to human sensitivity usually emerging in early layers—though for some high-level objectives the best- aligned layer shifted toward mid- to late layers (Flachot & Gegenfurtner, 2021; *Figures S8 and S9*). This task-independence may be specific to color discrimination: other chromatic phenomena, such as color categories and the chromatic contrast sensitivity function, emerge only for particular tasks (Akbarinia, 2025; Akbarinia, Morgenstern, & Gegenfurtner, 2023). This contrast underscores that the orange-biased asymmetry is largely a statistics-driven property.

Two methodological points deserve comment. First, stimuli were equiluminant with the background for the standard observer and were not adjusted for individual isoluminance, so the discs may have carried a small luminance signal. This cannot explain the hue–chroma asymmetry: by construction, hue and chroma shifts of a given magnitude produced equal changes along the L–M axis at each reference, so any luminance artifact would be of equal size for the two directions. Second, our design deliberately focused on two reference colors, orange and purple, where the hue–chroma asymmetry is most pronounced (Krauskopf & Gegenfurtner, 1992; Danilova & Mollon, 2016; Hedjar et al., 2025a, 2025b). Concentrating on these two colors allowed us to prioritize a comprehensive comparison of the same stimuli across environmental statistics, psychophysics, MEG with individually tailored stimuli, and neural networks; our conclusions are accordingly restricted to this contrast rather than to the geometry of color space as a whole.

Together, our results reveal a consistent pattern across levels of analysis. Environmental color statistics parallel the hue–chroma asymmetry of human color discrimination; the same asymmetry is reflected in task-dependent cortical activity and emerges spontaneously in artificial systems optimized for natural visual tasks. Most importantly, reversing the chromatic statistics of the training environment in artificial neural networks was sufficient to reverse the asymmetry itself. These findings suggest that the geometry of color discrimination is not solely determined by fixed sensory architecture, but is shaped through learning and adaptation to the structure of the visual environment. Ecological chromatic structure determines which color distinctions are represented with high resolution, whereas task-dependent cortical dynamics determine when those distinctions become expressed in neural activity and behavior.

## Methods

### Definition of DKL space

#### Psychophysical and physiological experiments

Colors were specified on the isoluminant plane in DKL color space, whose cone-opponent dimensions follow the activations of large groups of retinal ganglion cells in the eye and neurons in the LGN. We defined the space in terms of the projector output at midgray. We used a SpectroCAL MKII Spectroradiometer (Cambridge Research Systems) to measure the spectrum of the R, G, and B channels of the projector on the projection screen and converted the spectra to *xyY* values using Judd-corrected color matching functions. The DKL-to-RGB transformation matrix was:

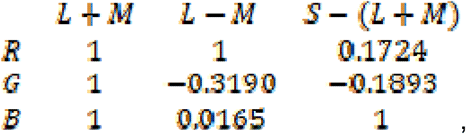

calculated using Smith and Pokorny (1975) cone fundamentals (with Boynton’s (1979) *Z* coefficient value). The CIE1931 (Judd-corrected) *xyY* values at midgray of the projection onto the screen were [0.339, 0.353, 49.1].

In DKL color space, the relationship between the scales of the S axis and the L-M axis is not explicitly specified, so it is convention to normalize them based on detection thresholds at the adaptation point, such that the thresholds are circular (i.e., a radius of one detection threshold unit; Krauskopf & Gegenfurtner, 1992). Typically, thresholds along the S axis are much larger than along the L-M axis, requiring us to scale down the S-axis values by a factor. To calculate this factor, we collected detection thresholds from six naïve participants (average age = 32.8, SD = 7.4, four females; five later completed the main psychophysical experiment) using a four- alternative forced-choice task. A uniformly colored 1° disc could appear in one of four locations around the center of the screen. The four locations were the same as specified in the diamond stimulus configuration of the psychophysics and MEG experiments, except that the diamond configuration was never tilted (see *Psychophysics-only experiment* below). The disc color differed from the background in the chroma direction along one of eight hue angles (although only the cardinal axes were analyzed). We predefined 15 chroma shifts along each hue angle, ranging from 0 to 0.08 DKL units away from the background. An adaptive staircase (QUEST; Watson & Pelli, 1983) determined the chroma shift for each trial. We initialized the prior threshold estimate and its standard deviation to 0.5 and 0.2, and set the threshold criterion to 0.5. Weibull parameters *beta*, *delta*, and *gamma* were set to 3.5, 0.01, 0.25, respectively.

Staircases across the eight hues were interleaved. We measured 60 trials per staircase and fitted the results using the *psignifit* software to a sigmoidal psychometric curve; thresholds were determined at 62.5% accuracy. As expected, thresholds were elongated along the S-axis. Based on our measurements, we used 2.53 as the scaling factor.

Stimuli were presented using a PROPixx VPX-PRO-5050B projector (VPixx Technologies) at 25% power, which has a resolution of 1920×1080 (52.2° × 29.0°), 8 bits/channel, and a linear RGB output. The projector was placed outside the magnetically shielded recording booth; stimuli were back-projected onto a semi-transparent projection screen placed at 1 m viewing distance. *xyY* coordinates of the RGB channels were: R = (0.613, 0.349, 20.3), G = (0.283, 0.605, 64.1), B = (0.157, 0.0709, 8.63). We used Psychtoolbox-3 on MATLAB R2018 as our presentation software.

### Psychophysics-only experiment

#### Stimuli

We defined two reference points in DKL space which bisected the orange (positive L-M and negative S-LM) and purple (positive L-M and positive S-LM) quadrants 0.2127 units away from the origin. The *xy* of the purple and orange references were [0.338 0.266 49.11] and [0.445 0.442 49.11], respectively. The discs were equiluminant to the background, a midgray color set to the origin of the DKL space (*xy* = [0.339 0.353]) at 49.11 cd/m^2^. As in previous work (Hedjar et al., 2025a, 2025b), we defined increasing and decreasing chroma as a shift towards and away from the adaptation point, and clockwise and counterclockwise hue as a shift along the tangent of the hue circle at the reference points. This ensures that one unit change in any direction asserts equal changes along the cardinal chromatic dimensions. We predefined 10 possible shifts in each of these directions, ranging from 0 to 0.06 units away from the reference point.

Four 1°-diameter discs lay in a 2×2 diamond configuration. The distance from top edge to bottom edge of the diamond was 3°. The configuration was randomly tilted ±10° on every trial. The tilt is only relevant for the orientation discrimination task performed during MEG data collection for the task-comparison experiment (see *Magnetoencephalography (MEG) experiment*), but we wanted to keep the same possible stimulus configurations for all tasks, so the diamond exhibited a tilt across all trials and experiments. Three of the discs always had the reference color (orange or purple) and the color of one disc (the odd one out; randomly positioned) was shifted from the reference in either hue or chroma by one of the predefined shifts. The discs lay on a uniform background set to the midgray of the projector.

#### Procedure

The experimental procedure resembled those of earlier experiments (Hedjar et al., 2025a, 2025b). We presented each stimulus for 500 ms and participants indicated using a button box which of the four stimuli was the “odd one out” in terms of color only. They were asked to respond after the stimulus disappeared, but we accepted responses any time after stimulus onset. Participants had unlimited time to give a response. They were given auditory feedback on their performance immediately after (click sound = correct; white noise = incorrect). After the feedback, there was an intertrial interval of 1000–1200 (jittered) ms before the next trial.

The hue or chroma shift of the odd one out was determined by the adaptive staircase method QUEST (Watson & Pelli, 1983); parameters were the same as in the S-axis-scaling experiment, except that the prior threshold estimate was initialized to 0.6. Participants completed 60 trials for each condition’s staircase for a total of 480 trials: 2 quadrants × 4 directions (2 hue, 2 chroma). All conditions were interleaved. Participants completed the trials in two 12-minute blocks, between which they could take a self-paced break. Both blocks started with 35 discard trials to allow participants to get used to the procedure and to adapt to the gray background. The first few of these trials were presented for longer durations, gradually shortening to 500 ms for the last few trials. These trials merged seamlessly with the experimental trials and were not used in analyses.

During the experiment, a fixation dot was always present on the screen. We asked participants to keep their eyes on the fixation dot during stimulus presentation. Blinking was allowed during the intertrial interval. Participants performed this experiment in the MEG chamber at 1 meter distance from the projection screen.

#### Analysis of discrimination thresholds

We used *psignifit* toolbox (Schütt et al., 2016) to fit responses to a sigmoidal psychometric curve with chance-level at 25% and the lapse rate set to 0%; we determined thresholds at 62.5% accuracy. To get a single threshold for each quadrant, color direction, and participant, we averaged the two measured thresholds (e.g., increasing and decreasing chroma). These thresholds were used to determine the colors for the MEG experiments.

After MEG data collection (see below), we recomputed thresholds using discrimination data from all experiments in order to obtain more precise estimates of discriminability. From this aggregated data we converted thresholds to sensitivities by taking the reciprocal (1/threshold) and computed the log_10_ of the hue-to-chroma sensitivities for each participant and quadrant (*Figure 3*). This local comparison between hue and chroma sensitivities asks whether discrimination is preferentially enhanced along the hue direction, rather than reflecting overall sensitivity to chromatic variation within each color region.

#### Participants

Forty-four naïve participants (29 female) with an average age of 26.0 (standard deviation = 5.2) completed the psychophysics-only task. One participant completed the task twice. Five of these participants also completed the S-axis scaling detection experiment. All participants gave informed consent and reported normal color vision. Visual acuity was normal or corrected to normal using MEG-compatible glasses.

### Magnetoencephalography (MEG) experiment

#### Stimuli

Stimulus presentation for the MEG sessions was the same as during the psychophysics-only session. However, instead of adapting the color of the odd disc via a staircase procedure, we presented a fixed subset of odd disc colors; reference disc colors remained the same. These colors were individually calculated based on each participant’s thresholds, computed after completing the psychophysics-only experiment. Each set was composed of 26 colors: 3 chromatic distances (small, medium, and large) from the reference for all 8 directions (increasing and decreasing chroma and clockwise and counter-clockwise hue for the purple and orange DKL quadrants), plus two colors equal to the reference. For each shift magnitude (e.g., small shifts), we ensured that the chromatically shifted colors were equidistant from their respective reference across all color directions, such that the activation difference along each cardinal mechanism was identical for every color-reference pair.

To show the difference in discrimination ability across the color directions despite equal activation, we defined the odd disc colors across small, medium, and large shifts to coarsely sample the psychometric curve for some color directions but not for others. In order to simplify the possible colors to choose, we grouped color directions based on thresholds. We applied a *k*- means clustering across the thresholds of the four color directions (orange and purple hue and chroma, averaged between the two polarities of each color direction) from each of the 44 participants, using squared Euclidean distance as a distance metric and running 100 replicates. Orange hue thresholds formed one cluster (orange-hue condition) and purple hue and chroma along with orange chroma thresholds formed the other (mixed conditions) (mean silhouette value = 0.74 on aggregate thresholds).

We computed the set of odd disc colors to coarsely sample the psychometric curves of the mixed conditions while falling above threshold for the orange-hue condition. Odd disc colors with no shift and with a small shift were sub-threshold. We intended for the medium and large shifts to be suprathreshold, but projector gamut limitations prevented us from achieving such values for all participants; therefore, colors with the large shift were as suprathreshold as possible and colors with the medium shift thresholds lay in between the small and large. The relationship between odd disc colors and participants’ psychometric curves (refitted from all collected data) are detailed in the *SI*.

We ran two additional sessions to compare the effect of the task on MEG signals. In both sessions of this task-comparison experiment, the set of odd disc color shifts straddled orange- hue thresholds while falling below threshold for the mixed conditions.

#### Procedure

The procedure across all sessions was similar to the psychophysics-only experiment with a few exceptions. The odd disc location was counter-balanced across locations (21-22 trials per location, per unique odd disc color). The jittered intertrial interval began immediately after stimulus offset. Responses were accepted until the end of the intertrial interval, after which auditory feedback was presented and the next trial began. Presentation of conditions was randomized across the session. Each session was broken up into 12 blocks, in between which participants could take a short break. Data collection took approximately one hour per session.

In the orientation discrimination session, participants indicated whether the diamond configuration of the stimulus was tilted more to the left or to the right. The diamond orientation was randomized on each trial.

Participants could easily perform the orientation task (96% correct across participants and odd disc conditions). Reaction times on average were 473 ms after stimulus onset and no significant difference was found between odd disc conditions (reaction times collapsed across participants: *F*(25, 30756) = 1.25, *p* = 0.1818). For the color discrimination task, performance depended on the odd disc conditions, as expected. Reaction times were higher than for the orientation task, on average 778 ms.

#### Participants

Twenty-nine of the 44 participants who completed the psychophysics-only task participated in the main MEG experiment (17 female; average age = 26.4, standard deviation of age = 5.8).

Thirty of the 44 participants who completed the psychophysics-only task participated in the task- comparison MEG experiments. We excluded three participants from the color discrimination task due to invalid data, leaving 27 (20 female; average age = 26.1, standard deviation of age = 6.0). Nineteen of those 30 participants completed the orientation discrimination task (14 female; average age = 26.6, standard deviation of age = 4.9).

#### MEG data recording

The MEG signal was recorded continuously at a sampling rate of 1000 Hz inside a μ-metal– shielded recording room (Vacuumschmelze, Hanau, Germany) using an Elekta Neuromag TRIUX triple sensor system with 102 magnetometers (Elekta Neuromag Oy, Helsinki). Vertical and horizontal eye movements were monitored simultaneously (Synamps amplifier; Neuroscan) by recording the electro-oculogram (EOG) from bipolar electrode placements at the outer canthi of both eyes (horizontal EOG) as well as from a unipolar placement below the right eye (vertical EOG). MEG and EOG were band-pass filtered online in the analog domain (0.03 to 330 Hz) and then subjected to Maxwell filtering (MaxFilter Software, Elekta Instrument AB, Stockholm, Sweden) for the elimination of spatial and spatiotemporal interferences via spatiotemporal signal space separation and to correct for head movements during the measurement. The MEG data were then down-sampled to, and digitized at, a sampling frequency of 500 Hz.

#### Epoching, artifact rejection, event-related fields

MEG data were analyzed offline using the FieldTrip toolbox in MATLAB (Oostenveld et al., 2011). The continuous data were epoched relative to the onset of the colored stimulus from -0.5 to 1.5 s, demeaned, and line noise was attenuated using a discrete Fourier transform filter at 50 Hz and its harmonics (up to 150 Hz). A high-pass filter of 0.03 Hz and a low-pass filter of 100 Hz were applied. The epoched data were first visually inspected using the function *ft_databrowser*, followed by semi-automatic artifact detection applying a peak-to-peak amplitude threshold criterion. The latter was adjusted for each participant to eliminate corrupted trials, which resulted in a rejection rate of 5-15% of the epochs (min: 4.88%, max: 13.62%, mean: 8.76%). Artifact- cleaned trials were finally binned into experimental conditions (26 in total: one for each chroma and hue shift magnitude of orange and purple).

#### MEG-based decoding analyses

Decoding was performed on all 102 magnetometer channels. Preprocessing. Data were low- pass filtered at 30 Hz and down-sampled to 120 Hz (ft_resampledata). Data from all conditions were concatenated into a continuous dataset to perform normalization across the entire recording. Specifically, data were z-scored per channel across time and then segmented back into their original trial structure. Single-trial baseline correction was applied (−0.2 to −0.1 s). Data were converted into trial × channel × time matrices for subsequent decoding analyses.

#### Multivariate pattern analysis

Multivariate pattern analyses were conducted at the single participant level using custom MATLAB scripts and the MVPA-Light toolbox (Treder, 2020). To improve signal-to-noise ratio, time series were smoothed using a Gaussian kernel (= 3 time- samples) before decoding. Classification was performed using a linear discriminant analysis, yielding time-resolved estimates of class accuracies. To determine whether variations in hue or chroma could be distinguished from the reference condition in neural responses, we trained separate binary classifiers for each hue and chroma shift, where each shift and the reference condition were treated as the two classes, and evaluated how well the classifier could distinguish between them. Trial counts (after artifact rejection) were balanced across odd disc color conditions by random subsampling to match the minimum number of trials per condition. A k-fold cross-validation procedure (k = 5) was implemented and repeated 15 times using random partitioning of trials into folds (*mv_classify_across_time*). To increase signal-to-noise, in each repetition, the trials were randomly grouped into sets of 3 within each condition and averaged. The final performance metric reflects the average across repetitions.

#### Statistical validation

Decoding results were statistically evaluated using custom MATLAB scripts and the nonparametric cluster-based permutation test (Maris & Oostenveld, 2007) implemented in the FieldTrip toolbox (*ft_timelockstatistics*; Oostenveld et al., 2011). Accuracies were first averaged across polarities of each color direction (e.g., increasing and decreasing chroma) for each participant. The resulting decoding time courses were compared against chance level (0.5) using a dependent-samples t-statistic computed across participants at each time point. Temporally adjacent time points exceeding a one-sided cluster-forming threshold of p < 0.05 were grouped into clusters, and the sum of the t-values within each cluster (*maxsum*) served as the cluster-level statistic. Significance was assessed with a Monte Carlo permutation procedure (10,000 randomizations). In each randomization, the condition labels (decoding accuracy versus chance level) were randomly swapped within participants, which is equivalent to sign-flipping each participant’s decoding-minus-chance difference time course; the t-statistics were then recomputed at every time point, thresholded, and clustered as described above. Observed cluster-level statistics were compared against the resulting permutation distribution of the maximum cluster-level statistic (one-sided test for above-chance decoding, α = 0.05). Note that statistical analyses were performed on temporally smoothed decoding time courses (Gaussian kernel, σ = 3 time samples). Smoothing may increase the temporal extent of significant clusters, which should be considered when interpreting cluster onsets and offsets.

### Analysis of chromatic statistics of reflectance dataset and image databases

We analyzed the hue distribution of reflectance measurements of natural objects, large-scale image datasets, a smaller set of calibrated camera images, and hyperspectral images. For each image set, all pixels were included in the analysis except achromatic pixels with a DKL chroma value < 0.01. Color conversion followed the procedure described in the subsection “Psychophysical and physiological experiments” within the “Definition of DKL space” section. The histograms in Figure 2 are plotted in unscaled DKL coordinates, without the psychophysical S-axis scaling used in the psychophysical and MEG experiments, so as to characterize the datasets without a display- and observer-specific normalization. This choice does not affect the results: scaling multiplies the S coordinate of every pixel by a constant, which changes hue angles within a quadrant but moves no pixel across the cardinal axes, so the quadrant proportions reported in the Results are identical with and without scaling.

Our analysis began with a natural reflectance dataset aggregated from 6,476 collections which include reflectances from object and material categories such as bark, flowers, fruits, grass, human skin and hair, leaves, lichen, pelage, plants, rocks, stones, snow, soil, tree logs, vegetables, and vegetation. These reflectance sets were compiled from multiple research groups worldwide (Arnold et al., 2008; Regan et al., 1998; Sumner & Mollon, 2000; Brown, 1994; Krinov, 1947; Matsumoto et al., 2014; Tajima et al., 1999; Morimoto et al., 2022). All reflectance data were rendered under a D65 illuminant and subsequently converted to DKL color space.

Secondly, we collected large-scale RGB image corpora that dominate contemporary machine learning. We assumed nominal sRGB encoding and linearized the pixel values using a gamma value of 2.4 (International Electrotechnical Commission, 1998). We processed Places365 (1,803,460 images, Zhou et al., 2017), ImageNet (1,431,167 images; Deng et al., 2009), Tiny Taskonomy (366,784 images, Zamir et al., 2018), Microsoft COCO (65,616 images; Lin et al., 2015), and THINGS (26,107 images; Hebart et al., 2019). These corpora differ in origin and purpose: Places365 was built for large-scale scene recognition, spanning 365 scene categories sampled from across the web; ImageNet was designed as a hierarchical benchmark for large- scale object recognition with extensive annotation; Tiny Taskonomy comprises indoor-scene photographs annotated for a broad battery of visual tasks; COCO emphasized common objects in context with dense instance segmentation; and THINGS curated naturalistic object images for psychology, neuroscience, and computer vision research. All datasets are widely used benchmarks, assembled largely from web sources and crowd-sourced annotations— approaches that provide statistical power but also introduce uncontrolled color pipelines (white balance, tone curves, in-camera transforms) with unknown deviations from sRGB.

We then analyzed calibrated natural image datasets in which camera responses were measured and images were provided in device-independent coordinates: the McGill Calibrated Colour Image Database (1,381 images; Olmos & Kingdom, 2004), the Barcelona Calibrated Images Database (448 images; Vazquez-Corral et al., 2009; Párraga et al., 2009; Párraga et al., 2010), and the València IPL Calibrated Color Image Database (65 images; Laparra et al., 2012; Gutmann et al., 2014). These datasets reduce uncertainty about image acquisition pipelines and enable more reliable colorimetric inference.

Finally, we examined hyperspectral images of natural scenes collected by seven groups, drawing on one or more datasets per group when available. These included the BGU ICVL dataset (165 images; Arad & Ben-Shahar, 2016), the Harvard Real-World Hyperspectral Images (77 images; Chakrabarti & Zickler, 2011), two Tokyo Tech datasets (51 images; Monno et al., 2019), including a 59-band visible–NIR corpus spanning 420–1000 nm at 10 nm intervals, the Foster and Nascimento hyperspectral database (47 images; Nascimento et al., 2002; Foster et al., 2006; Foster et al., 2016; Nascimento et al., 2016), the Columbia CAVE repository (31 images, 31 bands, 400–700 nm; Yasuma et al., 2008), and the Granada UGR outdoor dataset (14 images; Eckhard et al., 2015). When pixel values represented surface reflectance, spectra were rendered under D65 before cone excitations were computed; when they represented radiance, the measured spectra were used directly. For datasets extending into the near- infrared, bands outside the visible range were excluded before conversion to cone excitations and DKL coordinates. Hyperspectral data enable computation of cone excitations directly from spectral information, eliminating ambiguities introduced by consumer RGB encodings.

### Estimation of chromatic discrimination thresholds from neural networks

We focused on ResNet architectures (He et al., 2016) for two main reasons. First, their repeated residual blocks create a consistent structure that makes it straightforward to extract and systematically compare activity across different processing stages. Second, identical ResNet variants have been widely adopted for a range of everyday vision tasks and trained with different image datasets. This provided models with the same underlying architecture but exposed to distinct task demands and training data, allowing us to examine how both factors jointly shape the color sensitivities encoded in the networks.

We evaluated ResNet50 trained on ImageNet for object recognition (Deng et al., 2009), in which the model assigns a single category to the entire image (e.g., "dog," "car," or "chair"). We also tested the same architectures trained on Places365 (Zhou et al., 2017) for scene categorization, which involves recognizing environments such as kitchens, classrooms, or beaches. To move beyond whole-image classification, we included two ResNet50 models trained on the COCO dataset (Lin et al., 2015): Faster R-CNN for object detection (Ren et al., 2016), which must both locate and label multiple objects in an image (e.g., detecting "a person" and "a bicycle" together), and Keypoint R-CNN for human pose estimation (Ding et al., 2020). Because the publicly available COCO-trained Faster R-CNN and Keypoint R-CNN models are both initialized from ImageNet-pretrained ResNet50 backbones, their representations would confound the influence of the COCO task with that of ImageNet pretraining. We therefore trained these two networks from scratch on COCO, without ImageNet pretraining. Finally, we tested a custom ResNet50 trained on a modified version of ImageNet in which the sign of the S−(L+M) channel was inverted across all pixels, making purple hues the most common.

From each network, we recorded activity at six processing stages: the initial stage after the first pooling operation, the four successive residual blocks, and, when available, the final fully- connected layer before the output layer. Note that some network releases omit the final layer, expecting users to supply their own optimization. In all cases, the networks were kept frozen and not updated during our analyses.

To estimate chromatic discrimination thresholds from these representations, we attached a shallow readout classifier to each processing stage and trained it on a four-alternative forced- choice "odd-one-out" task. On each trial, the network was shown four simple 2-D objects that differed in shape or viewpoint. We used colored shapes rather than uniform discs, following Flachot and Gegenfurtner (2021), who varied the color of objects to probe color responses in deeper network layers. Uniform color fields may elicit less reliable responses in deeper layers, whereas colored shapes provide spatial structure that can drive activations throughout the network. The shapes used for threshold estimation were never seen during readout training. Three of them (the distractors) shared one RGB color, and the fourth (the target) had a different RGB color. The two colors were drawn independently and uniformly from RGB space, and the target’s position among the four was randomized. We then extracted the network’s frozen features for each of the four objects and passed them to the readout, which compared the four feature vectors using both their deviations from the four-object mean and the absolute size of those deviations, before a trainable linear layer selected which object was the odd one out. Because only the readout was trained while the network itself stayed fixed, any differences in the resulting thresholds reflected biases already present in the network’s representation rather than additional learning by the network.

After training the readouts, we estimated chromatic discrimination thresholds from classifier performance on held-out validation shapes using the same QUEST adaptive staircase framework used in the human psychophysical experiment (Watson & Pelli, 1983). Thresholds were measured around the orange and purple reference chromaticities on the isoluminant plane, separately along chroma and hue directions. This allowed direct comparison between human discrimination asymmetries and the chromatic information available at different depths of each network.

To manipulate the chromatic statistics of the training environment, we created a hue-inverted version of ImageNet by converting each pixel from linearized sRGB to DKL coordinates, inverting the sign of its S–(L+M) coordinate, and converting back to sRGB. This reflects every hue about the L–M axis, exchanging the frequencies of orange and purple (and of blue and green) while leaving luminance and chroma unchanged. Pixels that fell outside the sRGB gamut after inversion (8.62% of all pixels across the 1.28 million training images) were pulled radially back to the gamut boundary, which preserves hue; the median affected pixel was shortened to 94% of its original DKL vector length (interquartile range 83–98%), with a mean chroma loss of 0.050 DKL units and essentially no change in luminance (mean −0.005). A ResNet50 was then trained from scratch on this image set with the same architecture and training procedure as the standard ImageNet model, and thresholds were estimated as described above.

Full details of the stimulus generation, readout training procedure, network stages, optimization settings, and threshold estimation are provided in the *SI* (*Figure S11*).

## Acknowledgments

This project was funded by European Research Council ERC AdG Color 3.0 (884116) and by DFG Excellence Cluster EXC 3066/1 “The Adaptive Mind” (533717223). TM was supported by a Sir Henry Wellcome Postdoctoral Fellowship from the Wellcome Trust (218657/Z/19/Z). MVB and J-MH were supported by the German Research Foundation grant CRC1436 TP-B05 (Project ID 425899996). AA was supported by the Hessian Ministry of Higher Education, Research, Science and the Arts (LOEWE Start Professorship).

## Supplementary Information

### Individualized odd disc colors presented in MEG experiments

We tailored the odd disc colors presented during MEG data collection to each participant’s individual thresholds. *Figure S1* plots the chromatic distances in DKL units between each color shift and the orange or purple reference for each participant and each experiment. See *Methods* in the main text for an explanation of how we chose the colors for the two experiment sets.

**Figure S1.**
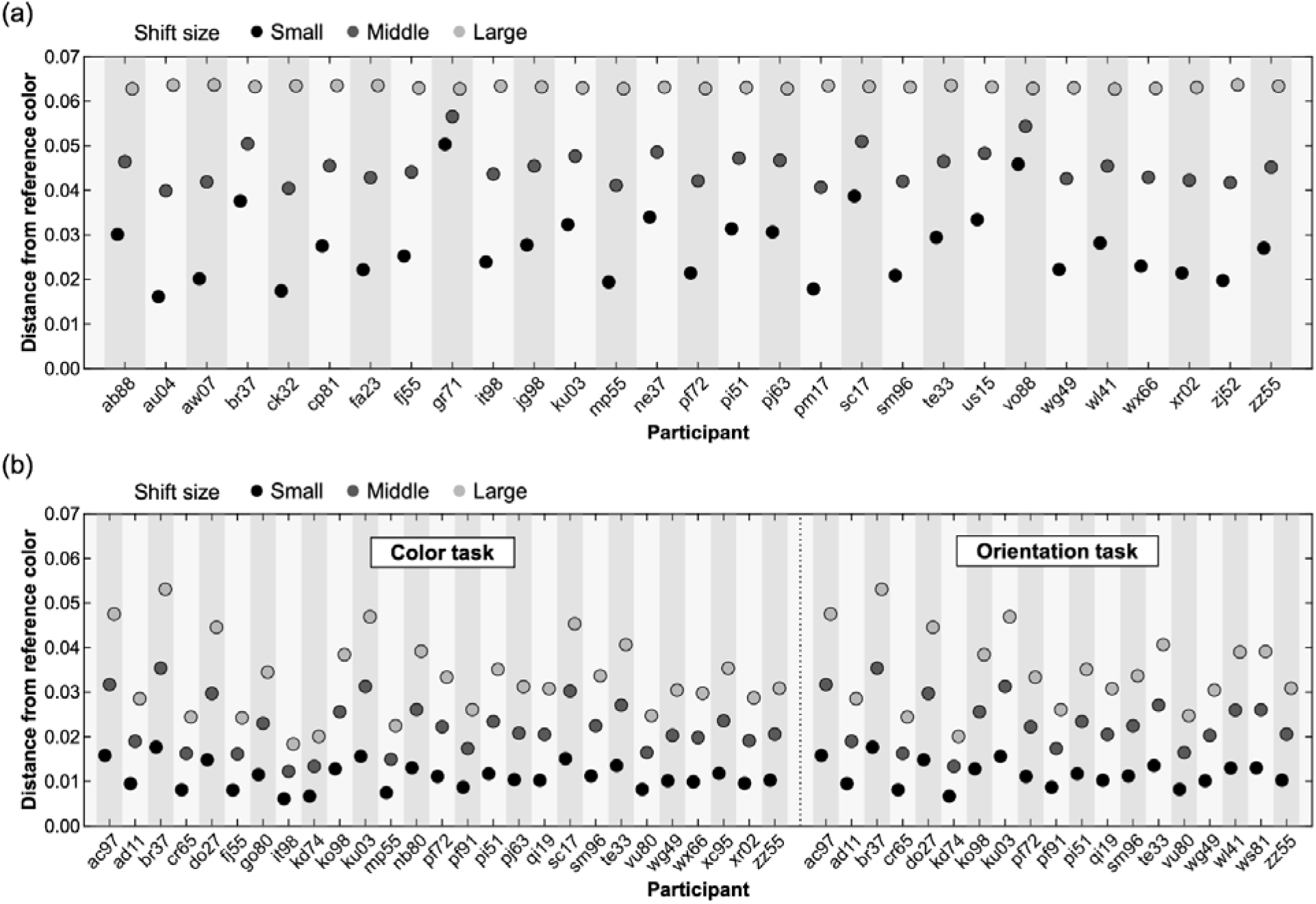
Distance between odd disc color shifts and reference colors used in MEG experiments. (a) Main MEG experiment. (b) Task-comparison experiments. For both panels, the y-axis plots the absolute distance between each color shift and the reference color (orange or purple) for each observer. The distance between reference and color shifts within a shift magnitude group (e.g., colors with a small shift from the reference) was always the same across color directions tested (i.e., orange increasing chroma, purple clockwise hue, etc.). All colors varied along both the S-(L+M) and the L-M dimensions (see Figure 3). Small, medium, and large color shifts are plotted as black, dark gray, and light gray dots.

Figure S2 plots the relationship between participant thresholds and the odd disc colors. Note that the set of colors for each participant was determined based on thresholds calculated from the psychophysics-only color discrimination data; however, here we fitted psychometric curves across responses given throughout all experiments in order to obtain more precise estimates of discriminability. For the main experiment, one can see that all odd disc color shifts along the orange hue direction lie almost entirely above the calculated thresholds of all participants. Along the other three directions, the color shifts span a wide range of the psychometric curve, from chance level to 100%. For the task-comparison experiments, we selected colors to produce the reverse pattern: the small, medium, and large shifts span the entire psychometric curve along the orange hue direction but fall almost entirely below thresholds for the other three directions.

**Figure S2.**
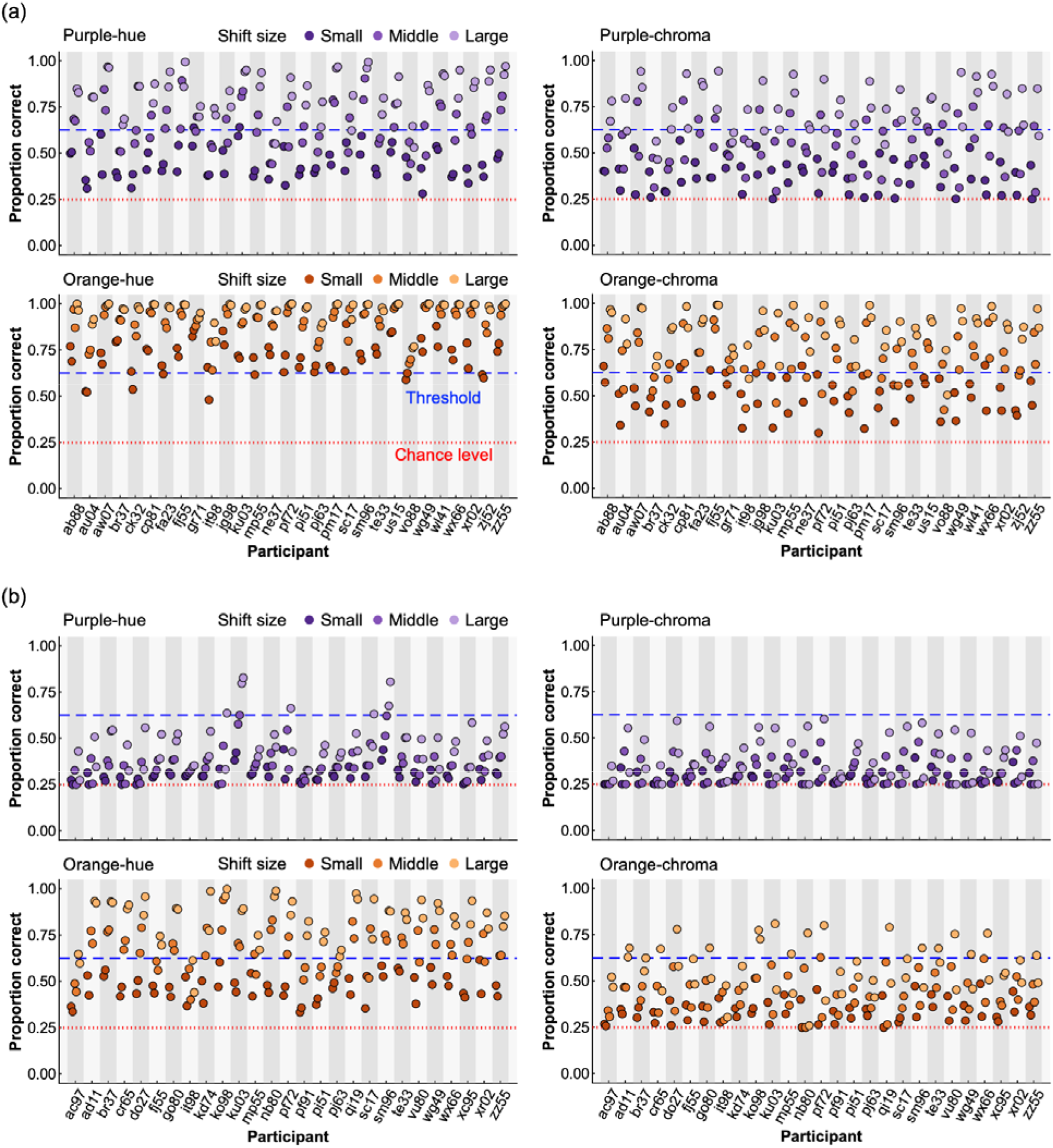
Fitted discrimination performances for odd disc color shifts used in MEG experiments. (a) Main MEG experiment. (b) Task-comparison experiments. Fitted proportion correct of each odd dis color shift per observer, color direction, and DKL quadrant. See Methods for details about the fitting of the psychometric curves. Odd disc colors were chosen using discrimination data from the psychophysics-only task, however we refit the curves from the discrimination data obtained across all experiments (psychophysics-only and both MEG color discrimination tasks) to compute the fitted performances shown here. We plot each color direction and quadrant separately. For each subplot, we show the fitted proportion correct of the small, medium, and large shifts (dark purple, purple, light purple dots for the purple quadrant; dark orange, orange, light orange dots for the orange quadrant). Because each direction has two polarities (increasing and decreasing chroma, clockwise and counter-clockwise hue), two dots are plotted side by side for every color shift.

#### Relationship between decoding accuracy, discrimination performance, and DKL color distance

To assess whether the decoding accuracy of the MEG signal was better explained by perceptual discriminability or physical chromatic distance, we related decoding accuracy to both fitted discrimination performance and absolute chromatic (DKL) distance between odd disc color and reference color. *Figure S3* plots both relationships across all data for the color discrimination tasks (main experiment and task-comparison experiment; panels *a* and *b*) and the orientation discrimination task (panels *c* and *d*). Across color discrimination tasks, discrimination performance correlated more highly with decoding accuracy (*r* = 0.654, *p* = 1.01×10^-164^) than did chromatic distance (*r* = 0.493, *p* = 3.33×10^-83^). In the orientation task, however, neither comparison showed a meaningful correlation (discrimination performance: *r* = 0.033, *p* = 0.4835; chromatic distance: *r* = -0.009, *p* = 0.8536).

To determine whether decoding accuracy was better explained by discrimination performance or by chromatic distance, we fitted linear mixed effects models with z-scored discrimination performance (zperf) and z-scored DKL distance (zdist) as fixed effects and by-participant random intercepts. We first asked whether discriminability explains variance beyond chromatic distance within the color discrimination tasks. The main and task-comparison color experiments were pooled, with experiment included as an additional fixed effect and predictors z-scored within this dataset (decoding ∼ zperf + zdist + experiment + (1|participant); n = 1,344 observations from 56 participants). Discrimination performance was a strong unique predictor of decoding accuracy (β = 0.054, 95% CI [0.049, 0.058], t(1340) = 21.5, p = 1.4×10⁻), whereas chromatic distance contributed no additional variance once discriminability was accounted for (β = 0.003, 95% CI [−0.003, 0.009], t(1340) = 1.07, p = 0.28). Decoding accuracy did not differ between the two color experiments (β = −0.004, 95% CI [−0.019, 0.011], p = 0.63). Although the two predictors were correlated as expected, the degree of collinearity was modest and did not compromise this partitioning (r = 0.71, variance inflation factor = 2.0).

We next asked whether these relationships differed between task types. Because comparing a significant effect in one task with a non-significant effect in another does not itself establish a difference, we fitted a single model to the pooled data in which task type (color vs orientation) interacted with both predictors, with predictors z-scored across the pooled dataset (decoding ∼ zperf × task type + zdist × task type + (1|participant); n = 1,800 observations from 75 participants). The effect of discrimination performance was significantly smaller in the orientation task than in the color tasks (interaction β = −0.052, 95% CI [−0.061, −0.044], t(1794) = −11.8, p = 6.1×10⁻³¹): the simple slope was 0.053 (95% CI [0.048, 0.057], p = 2.1×10⁻) in the color tasks but 0.000 (95% CI [−0.007, 0.008], p = 0.93) in the orientation task. The effect of chromatic distance was near zero in both tasks and did not differ between them (color: β = 0.004, 95% CI [−0.002, 0.009], p = 0.18; orientation: β = 0.004, 95% CI [−0.007, 0.014], p = 0.50; interaction β = 0.0001, 95% CI [−0.012, 0.012], p = 0.99). Overall decoding accuracy was also lower in the orientation task (β = −0.043, 95% CI [−0.056, −0.029], t(1794) = −6.24, p = 5.5×10⁻¹).

Because chromatic distance and discrimination performance are themselves correlated, these results do not imply that physical color distance is irrelevant; rather, they show that behavioral discriminability carried the explanatory power once both predictors were allowed to compete, and that it did so specifically in the tasks for which color was behaviorally relevant. Together, these analyses support the interpretation that variation in MEG color decoding accuracy primarily reflected how discriminable the chromatic differences were to participants, rather than the physical distance of the odd disc color from the reference in DKL space, and that chromatic information was not reliably expressed in the MEG signal when color was irrelevant to the behavioral judgement.

**Figure S3.**
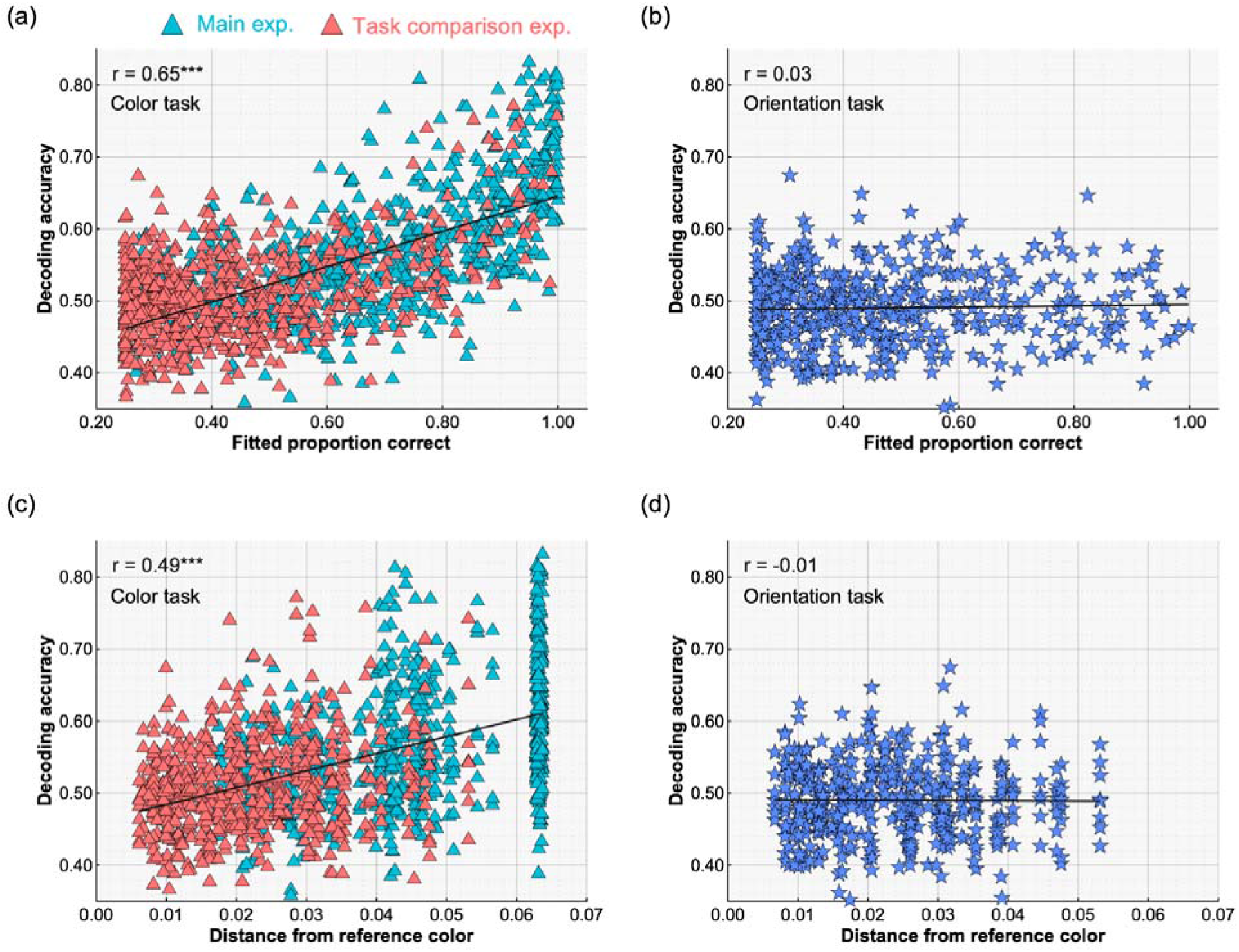
Relationship between fitted discrimination performance, absolute chromatic distance, and MEG decoding accuracy of the odd disc colors. MEG decoding accuracy, averaged between 350 and 650 ms after stimulus onset, is plotted against two measures of stimulus strength. (a, b) Decoding accuracy as a function of fitted discrimination performance (proportion correct). (c, d) Decoding accurac as a function of absolute chromatic distance from the reference color in DKL space. Panels (a) and (c) show decoding accuracies during the color discrimination tasks (turquoise = main experiment, red = tas comparison experiments). Panels (b) and (d) show decoding accuracies during the orientation discrimination task. Dots represent individual participant values for a single condition (e.g., purple clockwise hue, small shift). We fitted a line across points and computed Pearson’s r correlation coefficients.

### Individual participant patterns of MEG-derived hue–chroma asymmetry across shift magnitudes and task conditions

We plotted hue-to-chroma ratios from psychophysics and MEG, with the purple quadrant on the x-axis and the orange quadrant on the y-axis (*Figure S4a–c*) for individual participants. For psychophysics, the ratio was computed from discrimination sensitivity; for MEG, it was computed as the hue-to-chroma odds-ratio of decoding accuracy. MEG-derived ratios were computed separately for each participant and each odd disc color shift magnitude, using decoding accuracies averaged over 350–650 ms after stimulus onset.

Across the three color shift magnitudes, individual MEG participants showed a broadly similar qualitative structure to the psychophysical data. The MEG-derived points were generally displaced above the diagonal, indicating a stronger hue advantage for the orange color region than for the purple. This pattern was weak for the small shift, where individual participants showed substantial variability and the MEG group mean remained close to the origin. For the medium and large shifts, the MEG group mean shifted upward and moved closer to the psychophysical group mean, indicating that the MEG-derived neural asymmetry became more similar to the behavioral hue-superiority pattern as the stimulus shift magnitude increased.

This convergence was supported by participant-matched distance analyses. For each participant and MEG color shift magnitude, we computed the Euclidean distance in log-ratio space between the participant’s MEG-derived point and their own psychophysical point. These distances decreased across color shift magnitudes. MEG-derived points for the medium shift were significantly closer to psychophysics than those for the small shift (mean difference = 0.087, 95% CI [0.052, 0.123], t(28) = 5.03, p = 2.56 × 10⁻). Large shifts were also significantly closer than small shifts (mean difference = 0.114, 95% CI [0.085, 0.143], t(28) = 8.11, p = 7.83 × 10⁻). Large shifts were numerically closer than medium shifts, but this difference did not reach significance (mean difference = 0.026, 95% CI [−0.006, 0.059], t(28) = 1.66, p = 0.108). Thus, the individual participant MEG patterns became substantially more similar to behavior for medium and large color shifts, consistent with the group-level pattern shown in the main figure.

We also examined individual participant patterns in the task-comparison experiment (panel *d*). Participants performed a color discrimination task and an orientation discrimination task using matched visual stimuli. The color discrimination task produced a MEG-derived hue–chroma asymmetry that was closer to the psychophysical hue-superiority pattern, with a stronger asymmetry along the orange axis. Despite low average decoding performance for purple colors (Figure S6), many points lay to the right of the vertical zero line (which corresponds to a 1:1 hue-to-chroma ratio for purple), indicating numerically stronger purple hue decoding than purple chroma for many participants, although the difference was not significant (two-tailed one-sample *t*-test: mean log-ratio = 0.020, 95% *CI* [-3.65×10^-3^, 0.0437], *t*(26) = 1.74, *p*=0.0934). In contrast, the orientation discrimination task produced a weaker and less behaviorally aligned pattern (see main text for statistics).

Together, these individual-participant analyses show that the MEG-derived hue–chroma asymmetry was not a fixed property of the stimulus set alone. Instead, its alignment with behavior depended on both chromatic shift magnitude and task demands. Larger chromatic shifts produced neural decoding patterns that more closely resembled each participant’s behavioral asymmetry, and explicit attention to color enhanced the alignment between neural decoding and psychophysical hue superiority. This supports the interpretation that MEG decoding captures a task-sensitive cortical representation of the orange-specific hue advantage observed behaviorally.

**Figure S4.**
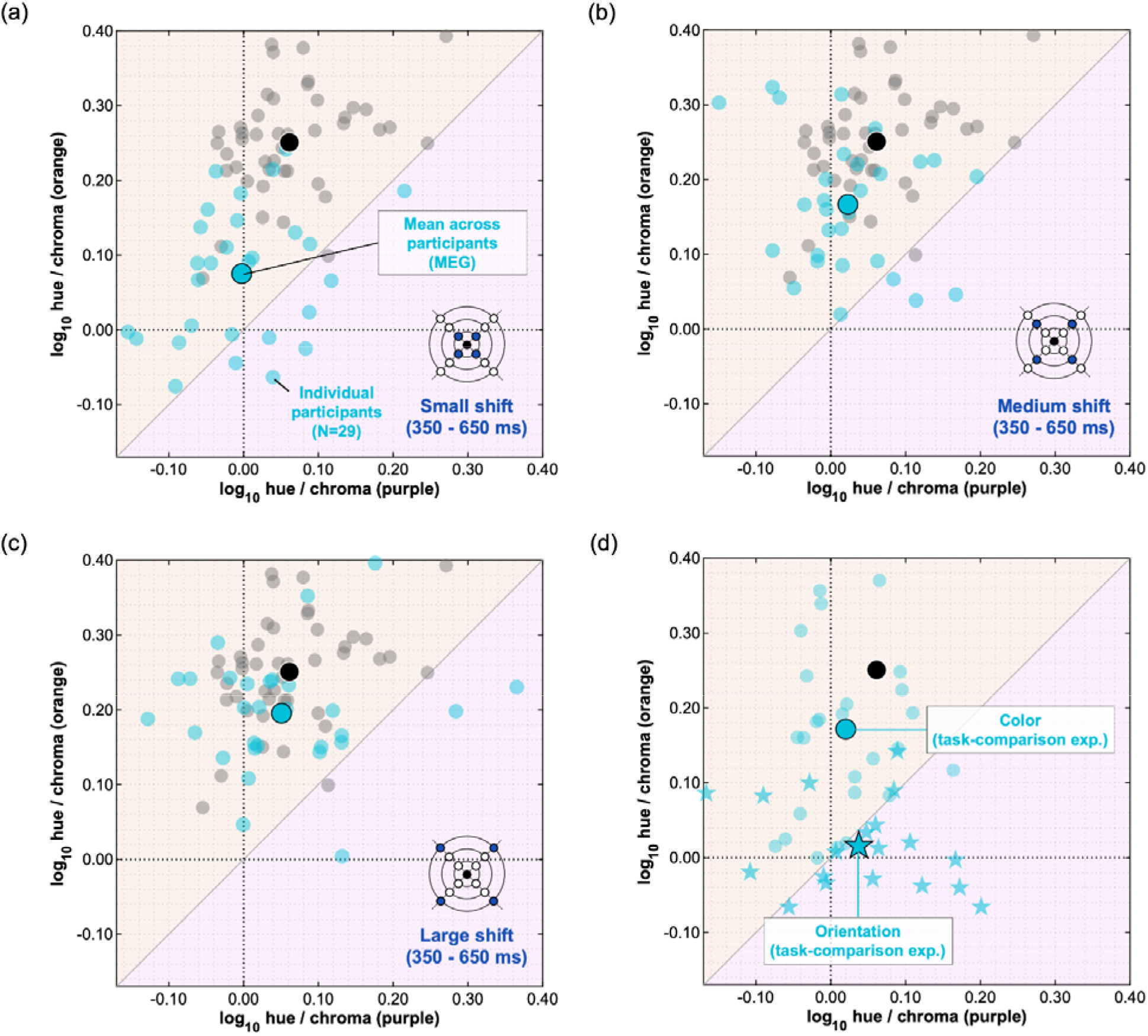
Relationship between psychophysical hue superiority and MEG-derived hue–chroma decoding asymmetries across color shift magnitudes and task conditions. (a–c) Hue-to-chroma log- odds-ratios are plotted for purple on the x-axis and orange on the y-axis. Gray circles show individual participants from the psychophysics-only experiment, and the black circle shows the psychophysical group mean. Cyan circles show individual MEG participants, and the cyan circle with a black outline shows the MEG group mean. MEG-derived log-odds-ratios were computed from decoding accuracies averaged over 350–650 ms after stimulus onset. Panels show results for the (a) small, (b) medium, and *(c) large odd disc color shift magnitudes. Across increasing shift magnitudes, the MEG group mean moved closer to the psychophysical group mean. (d) Hue-to-chroma log-odds-ratios of the large color shift from the task-comparison MEG experiments. Cyan circles show individual participants and the group mean for the color discrimination task, and cyan stars show individual participants and the group mean for the orientation discrimination task. The color discrimination task showed a hue–chroma asymmetry closer to the psychophysical hue-superiority pattern than the orientation discrimination task*.

### Cortical source activity elicited by colored discs during color discrimination

Source localization with MEG has been shown to reliably resolve activity in human visual cortex, including in our previous studies (Bartsch et al., 2015, 2021, 2023; Schulz et al., 2024). *Figure S5a* shows the source activity elicited by the colored disc stimuli during the main color discrimination task, across reference colors, color directions, and shift magnitudes. Each participant’s sensor data were co-registered with reference to the MNI-cortex (‘Standard Cortex’ in the source localization software Curry 8.3, Compumedics Neuroscience USA Ltd.) using individual anatomical landmarks (nasion, left and right pre-auricular point). A current source density (CSD) model was then estimated from the grand average (over participants) event- related magnetic field (ERMF) distribution at each time sample using the minimum norm least squares approach as implemented in Curry 8.3. An average cortical surface (‘Standard Cortex’) with 20,006 rotating dipoles served as a source compartment. Noise estimates were obtained from the baseline activity within a 500 ms period before stimulus onset. CSD estimates were then transformed to the FreeSurfer data format (*fsaverage*; FreeSurfer v7.4.1, https://surfer.nmr.mgh.harvard.edu/) using custom-made MATLAB scripts to allow us to overlay the borders of retinotopically defined visual areas V1, V2, hV4, PITd, and LO2 (CsurfMaps1; Sereno et al., 2022) onto the CSD maps using FreeView (v3). In this overlay, early visual cortex (EVC) areas V1 and V2, and mid-level ventral visual cortex (VVC) areas hV4, LO2, and PITd are highlighted. Note that this parcellation map reflects the average situation of cortical areas in the *fsaverage* surface. It is meant to give orientation regarding the approximate localization of source activity in the visual cortex. The source waves in *Figure S5b* illustrate the time course of activity within regions corresponding to the major source maxima. They were derived by averaging the CSD estimates at each time-sample across all current dipoles contained in the regions of interest.

The earliest source maximum appeared approximately 150 ms after stimulus onset in early visual cortex and adjacent ventral and lateral occipital cortex in the right hemisphere. Subsequent maxima emerged around 310 ms and 580 ms in ventral and lateral regions of the VVC (*Figure S5a*). Among those regions, hV4 is known to participate in coding of chromatic stimuli (Mullen et al., 2007; Bannert and Bartels, 2017, 2025; Schoenfeld et al., 2007), PITd in mediating endogenous attentional control of sensory selection (Stemmann and Freiwald, 2016, 2019), and LO2 being a part of a larger complex of lateral occipital areas in object processing (Kanwisher et al., 1996; Malach et al., 1995; Grill-Spector and Malach, 2004). Thus, source activity progressed from early visual areas toward a distributed ventral occipital network including regions associated with color processing and visual selection.

**Figure S5.**
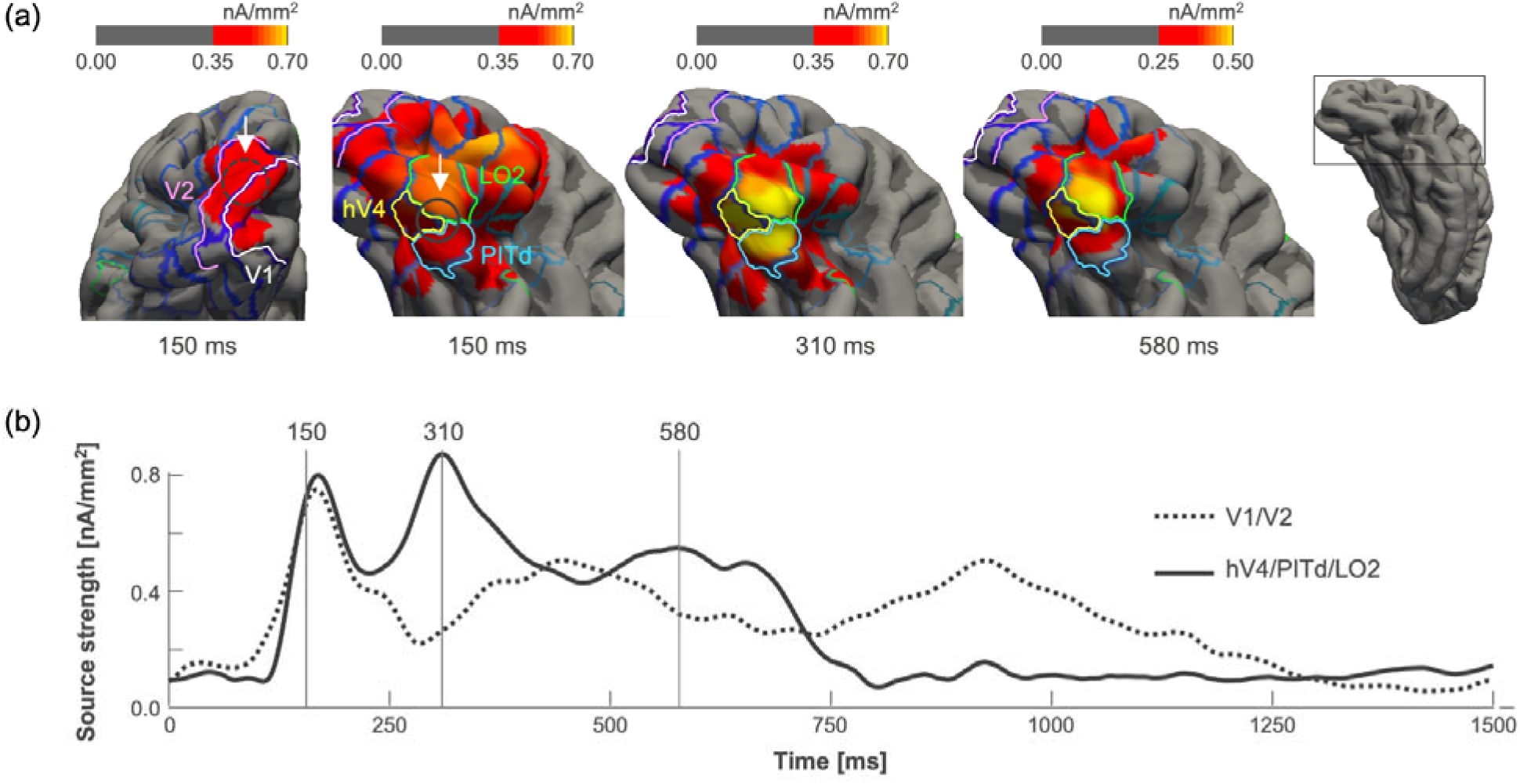
Source localization of the average magnetic field response elicited by the colored discs. (a) Topographical distribution maps of grand-average source activity (nanoamperes per square millimeter) estimated from the event-related magnetic field (ERMF) response using the minimum norm least squares approach as implemented in Curry 8.0.6 (Compumedics, Ltd.) at selected time points after stimulus onset across reference colors, odd disc color directions, and shift magnitudes. Activity is rendered onto the FreeSurfer average cortical surface (fsaverage) with a grand-average parcellation map overlaid (FreeSurfer’s CsurfMaps1; Sereno et al., 2022). Shown are ventral views onto the occipital part of the visual cortex. Retinotopically defined areas V1 and V2 are outlined in white and purple, respectively. Areas hV4, PITd, and LO2 are outlined in yellow, blue, and green, respectively. (b) Time- course of source activity (source waves) in regions of interest in early visual cortex and in mid-/high-level ventral extrastriate areas (encircled in dashed black and solid black lines in (a) and highlighted by white arrows). Time zero denotes stimulus onset, and stimuli disappeared after 500 ms.

### Task-dependent MEG decoding of chromatic differences

*Figures S6 and S7* plot the time-course of decoding accuracy for the task-comparison experiments (color discrimination and orientation discrimination).

**Figure S6.**
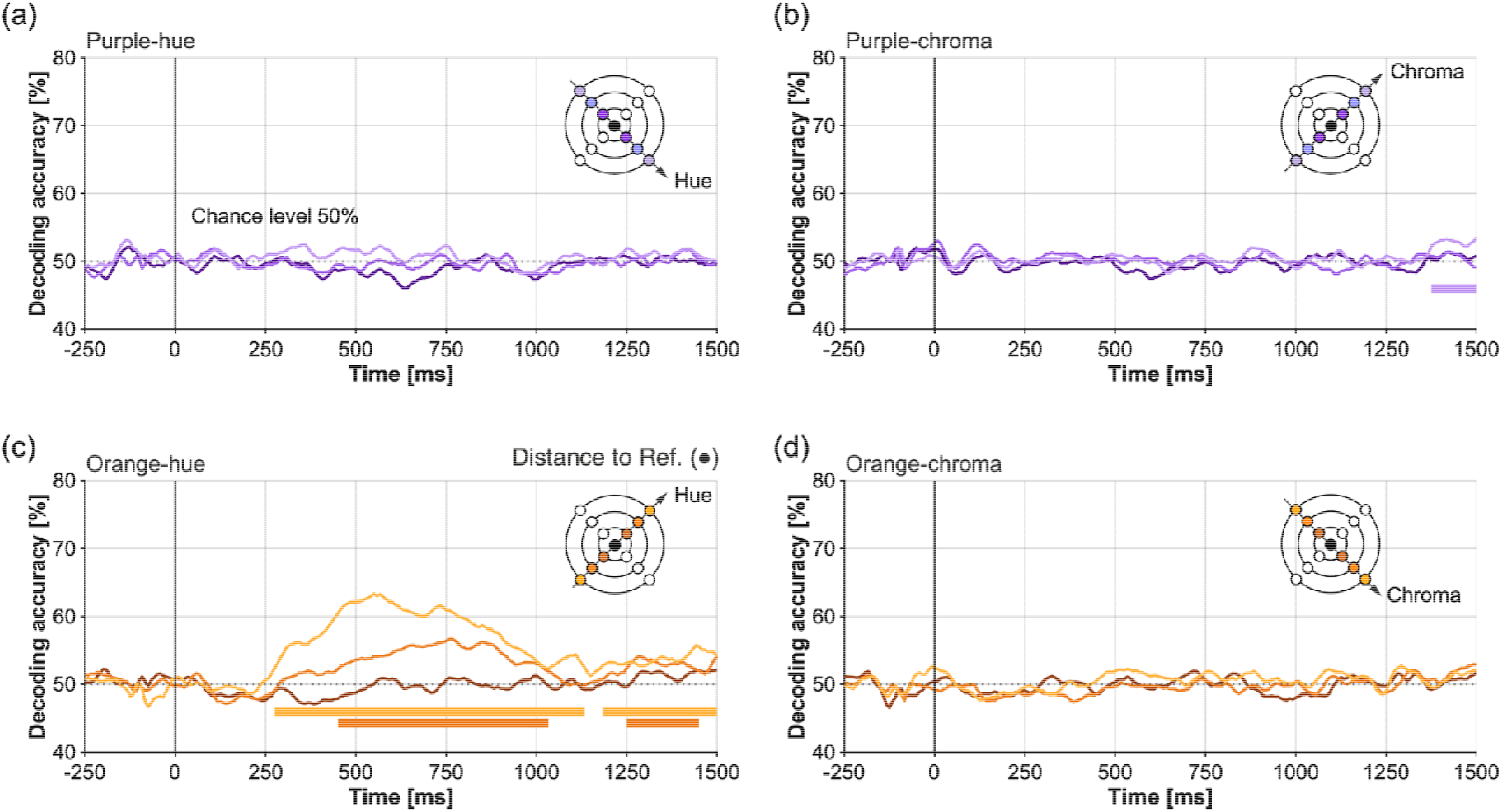
Multivariate pattern analysis of the odd disc color shifts for the color discrimination task in the task-comparison experiment. (a-b) Average decoding accuracy (% correct classification) for purple colors: small, medium, and large odd disc color shifts (plotted as dark purple, purple, and light purple lines, respectively) for (a) hue and (b) chroma. (c-d) Average decoding accuracy for orange colors: small, medium, and large odd disc color shifts (plotted as brown, orange, and light orange lines) for (c) hue and (d) chroma. Time zero denotes stimulus onset in panels (a-d), and stimuli were displayed for 500 ms. The horizontal bars indicate the time range of significant decoding (Monte Carlo significance probability p<.05) obtained with cluster-based permutation testing.

**Figure S7.**
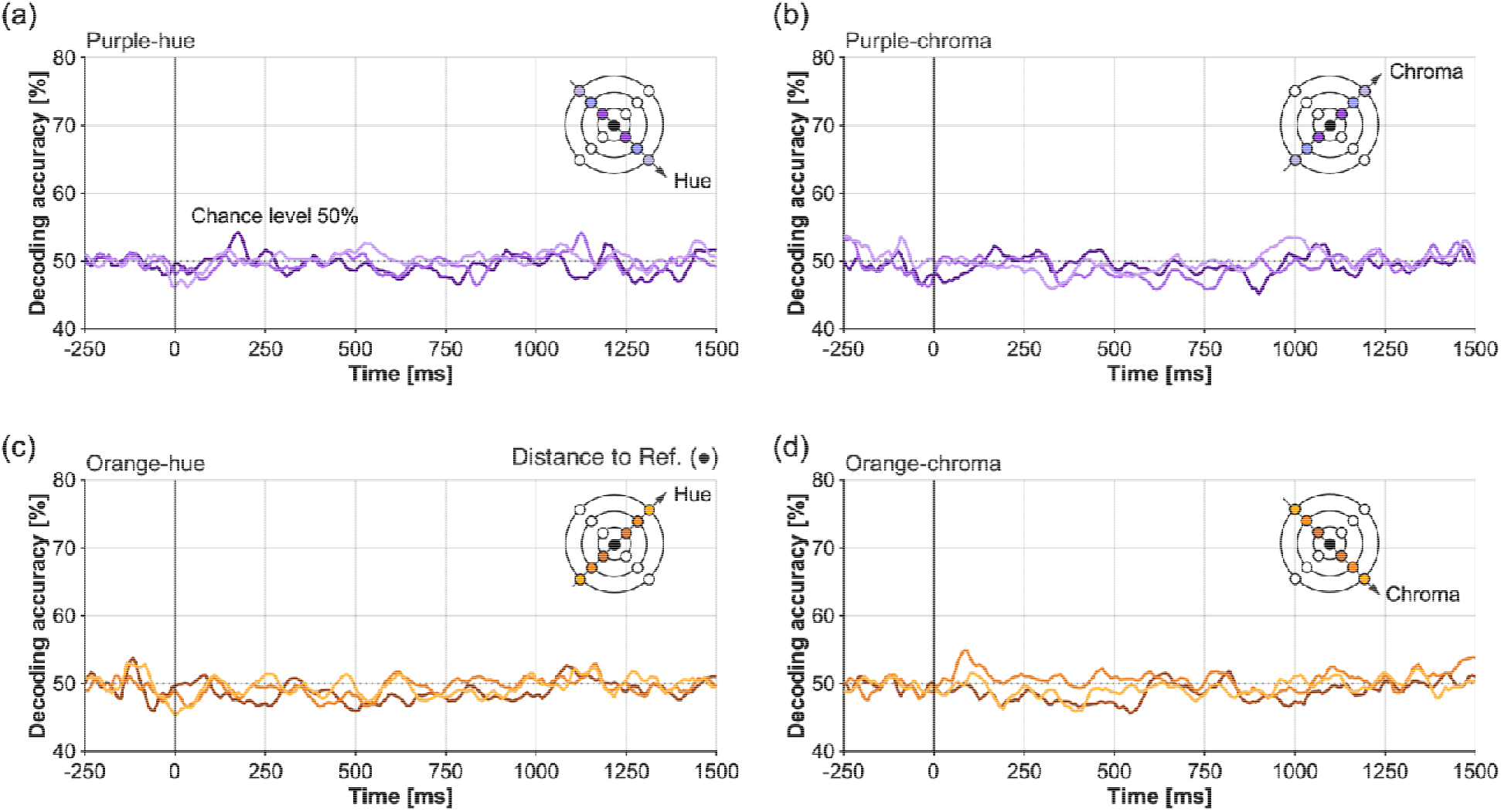
Multivariate pattern analysis of the odd disc color shifts for the orientation discrimination task in the task-comparison experiment. Panel layout follows that of Figure S6.

### Effects of task demands on color discrimination thresholds in deep neural networks

Here, we asked whether the human-like threshold pattern in the networks presented in the main text generalizes across a broader range of visual tasks, and whether task demands modulate the degree to which such an asymmetry emerges. The Taskonomy models (Zamir et al., 2018) are particularly informative in this respect, because they share an identical backbone architecture (ResNet50) and were trained on the same image database (Taskonomy including varied indoor scenes) but were optimized for different visual objectives. Taskonomy includes 25 tasks in total; however, we analyzed 24 of them, excluding the colorization task. The analyzed tasks and what each network learns to predict are summarized in Table S1. Colorization was excluded because the network operated on one-channel grayscale input images, which made it impossible to estimate chromatic thresholds.

For the color statistics analysis in *Figure 2*, we did not use the entire Taskonomy dataset, because it is extremely large (> 10 TB). Instead, we used "Tiny-Taskonomy," a smaller version of the same dataset that the dataset’s authors provide as a standard subset. It contains 35 of the 537 buildings (about 7%) and 366,784 images (about 8% of the full set). Because these images are drawn from the same pool of indoor scenes, we expect their color distribution to be broadly similar to those of the full dataset.

**Table S1.**
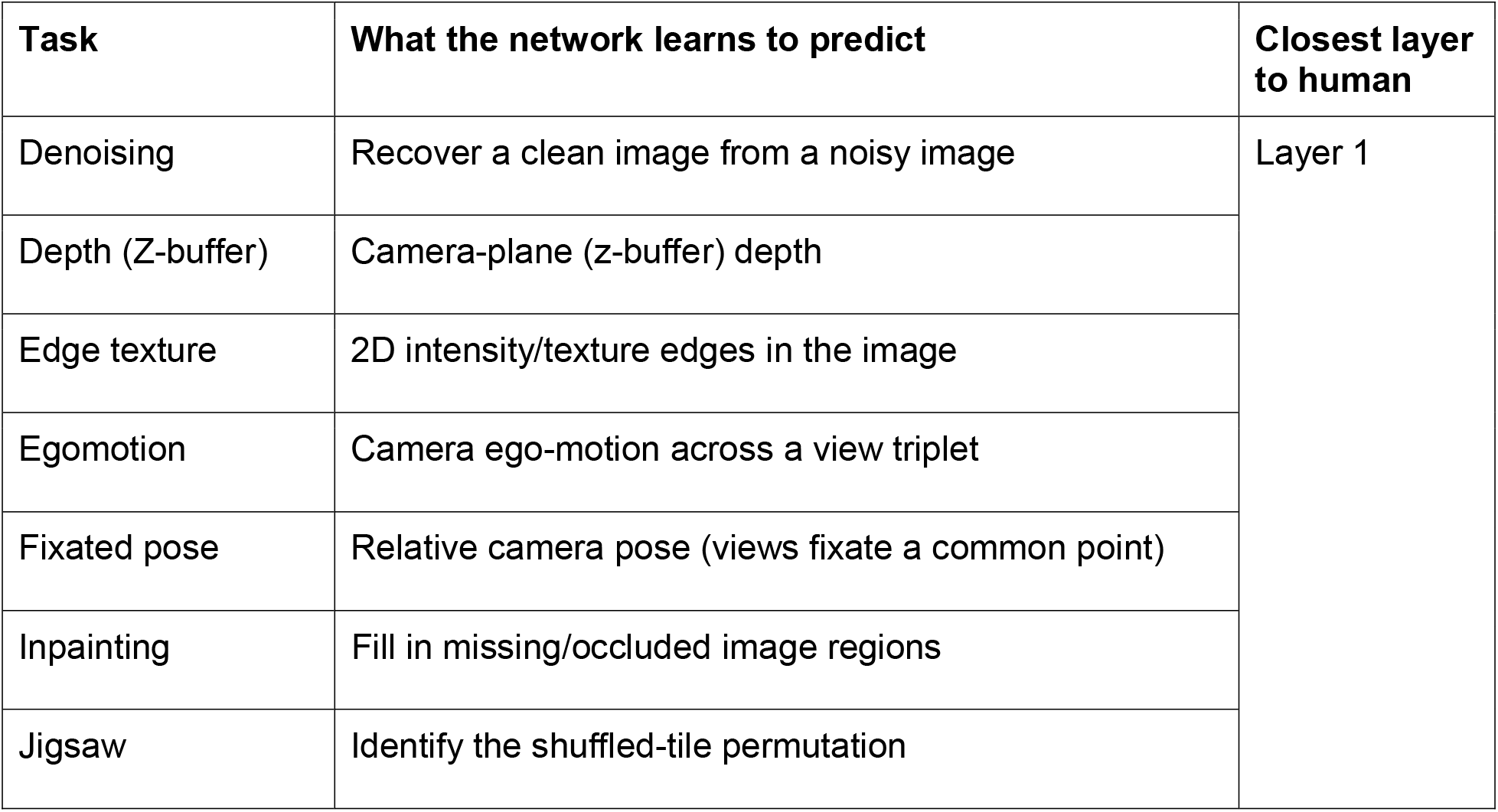
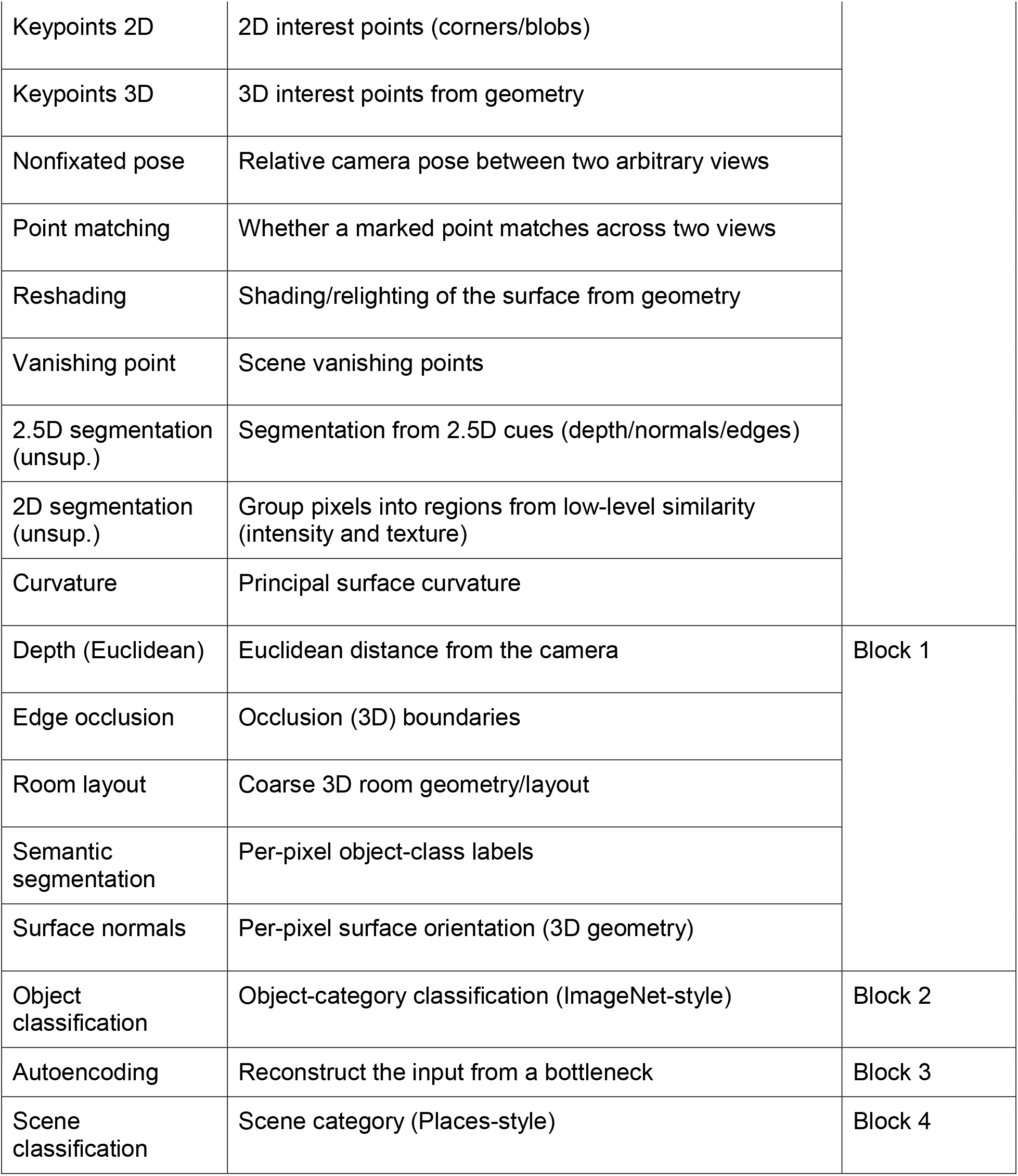
The 24 Taskonomy tasks and the layer that best aligned with human behaviors.

For each task-trained network we estimated hue and chroma discrimination thresholds at every processing depth (Layer 1, Block 1, Block 2, Block 3 and Block 4) and computed the log- transformed hue-to-chroma sensitivity ratio for the orange and purple references at each depth. To summarize how closely each network matched human perception, we identified, for every task, the single processing depth whose orange and purple sensitivity ratios lay closest to the human psychophysical mean (the best-aligned depth). *Figure S8* plots each task at this best- aligned depth, with marker color indicating which depth it was; Table S1 lists the best-aligned depth for every task.

Across the 24 Taskonomy-trained networks the human-like asymmetry was remarkably general: with a single exception, sensitivity ratios were larger for orange than for purple, placing the tasks above the diagonal in *Figure S8*. The orange-region asymmetry was therefore not restricted to the object recognition and scene classification networks discussed in the main text, but emerged across a wide range of visual objectives trained on the same natural image statistics. The only exception was unsupervised 2D segmentation, which fell below the diagonal. Where the human-like asymmetry was most closely matched, however, depended on the task. For 21 of the 24 networks the best-aligned depth fell at one of the two shallowest stages, Layer 1 (13 tasks) or Block 1 (8 tasks), and only three networks aligned best at deeper stages: object classification (Block 2), autoencoding (Block 3), and scene classification (Block 4). The tasks whose alignment peaked at the shallow Layer 1 and Block 1 stages were predominantly geometric, camera-geometry, and low-level image tasks—surface normals, curvature, Euclidean and Z-buffer depth, occlusion and texture edges, 2D and 3D keypoints, vanishing point, reshading, room layout, 2.5D segmentation, camera-pose and egomotion estimation, point matching, and the jigsaw and image-restoration (denoising and inpainting) objectives—for which color is presumably not directly diagnostic of the target output.

**Figure S8.**
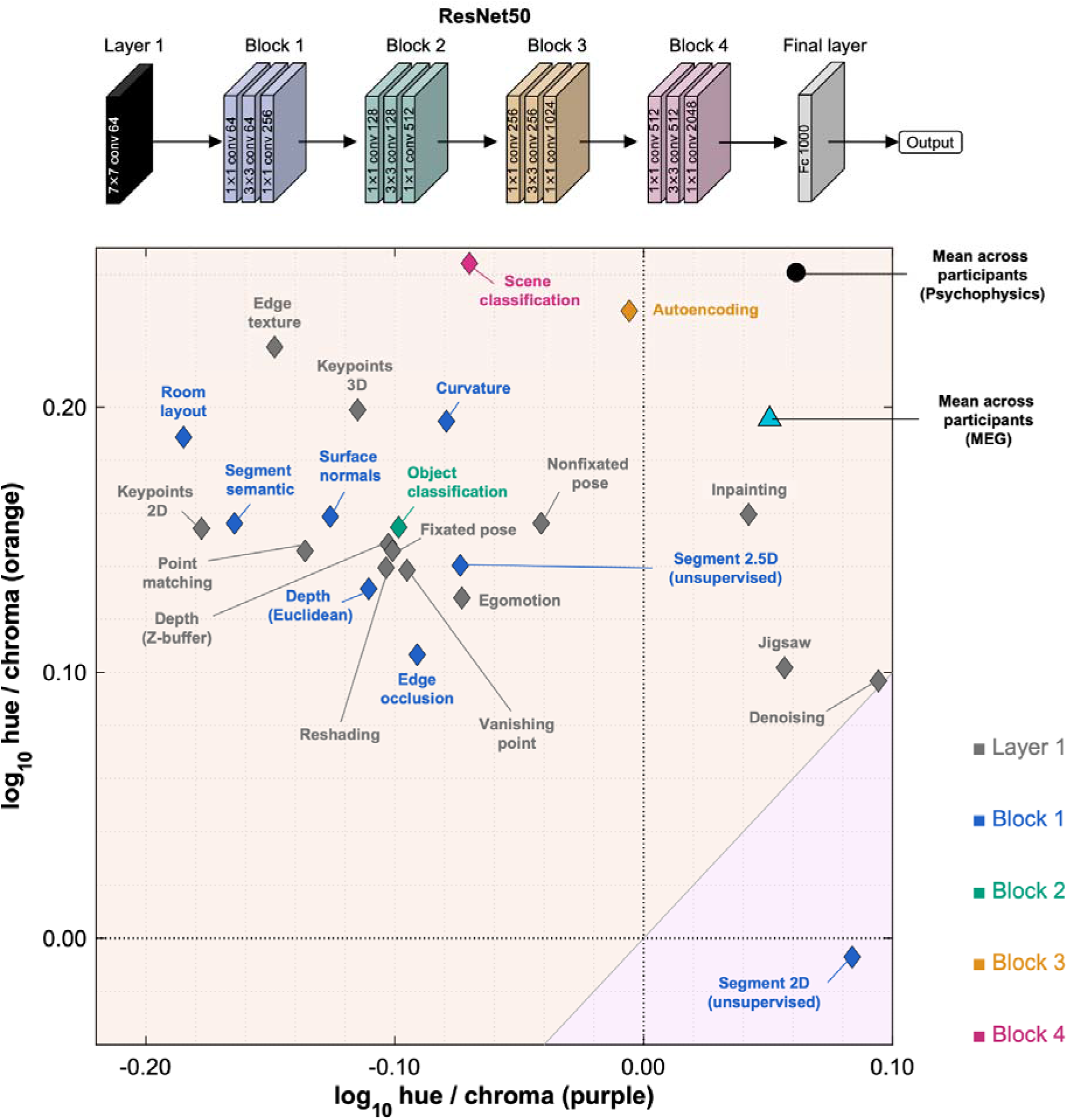
Hue-to-chroma sensitivity ratios across 24 Taskonomy tasks. Each symbol shows one ResNet50 network trained on a different Taskonomy task (Zamir et al., 2018), plotted by its log- transformed hue-to-chroma sensitivity ratio for the purple reference (x-axis) and the orange reference (y- axis). Positive values indicate higher sensitivity to hue than to chroma. For each task, the ratios are taken from the single processing depth whose values most closely match the human psychophysical mean (the best-aligned depth), and marker color denotes that depth (Layer 1, Block 1, Block 2, Block 3, or Block 4; see legend). Points above the diagonal show greater hue superiority in the orange region than in the purple region (the human-like pattern); all tasks except unsupervised 2D segmentation fall in this region. The black circle and cyan triangle show the group means of the human psychophysical and MEG data, respectively. The architecture, training database, and threshold-estimation procedure were identical across tasks, so differences reflect the training objective alone.

Examining the full depth profiles confirmed and refined this interpretation (*Figure S9*). The reconstruction and appearance tasks—autoencoding and inpainting, together with scene classification and the other 2D-appearance tasks (edge texture and 2D keypoints)—maintained a strong, comparatively flat orange-region hue superiority from early to late layers, consistent with the idea that these objectives encourage the preservation and use of color information throughout the network. By contrast, the geometric and 3D-structure tasks typically showed a high orange ratio at the early layers that declined toward deeper layers, often entering the human band only in the first one or two blocks before falling toward zero. Unsupervised 2D segmentation showed an orange ratio that became progressively more negative with depth while the purple ratio remained near zero, amounting to a depth-driven reversal of the typical asymmetry. This account is consistent with a general picture of how chromatic representations evolve with depth: irrespective of the eventual task, the early layers develop a generic, task- agnostic representation of color that closely reflects the chromatic statistics of the training images—which are themselves biased towards orange and blue (*Figure 2*)—and, as activity propagates to deeper, more task-specific stages, this representation is reshaped to serve the objective. When color is directly useful for the task, as in autoencoding (which must reconstruct the input image, including its chromatic content) and scene classification (in which diagnostic color regularities such as sky, vegetation, and ground remain informative), the orange-dominant asymmetry is preserved into the deeper layers; when color is not directly diagnostic, as in most geometric and 2D tasks, the asymmetry is gradually discarded, leaving the closest match to human sensitivity in the early, statistics-driven representations.

Taken together, these analyses point to a property that may be specific to color discrimination: the natural image statistics inherited during training appear to matter more than the particular visual task. This task independence is not a general feature of color phenomena in artificial networks: human-like color categories arise only in networks trained on high-level tasks (Akbarinia, 2025), and a human-like chromatic contrast sensitivity function only for a few objectives, most notably denoising and autoencoding (Akbarinia, Morgenstern, & Gegenfurtner, 2023). In a related vein, human subjective image-quality assessments align most strongly with a mid-level, V1-like layer, again implicating representations beyond the initial chromatic stage (Hernández-Cámara et al., 2025). One interpretation is that fine chromatic discrimination is governed largely by the low-level chromatic statistics of natural images, so it is established early and is relatively insensitive to task demands, whereas categorical and spatio-chromatic phenomena depend on the higher-level representations shaped by the task in deeper layers.

**Figure S9.**
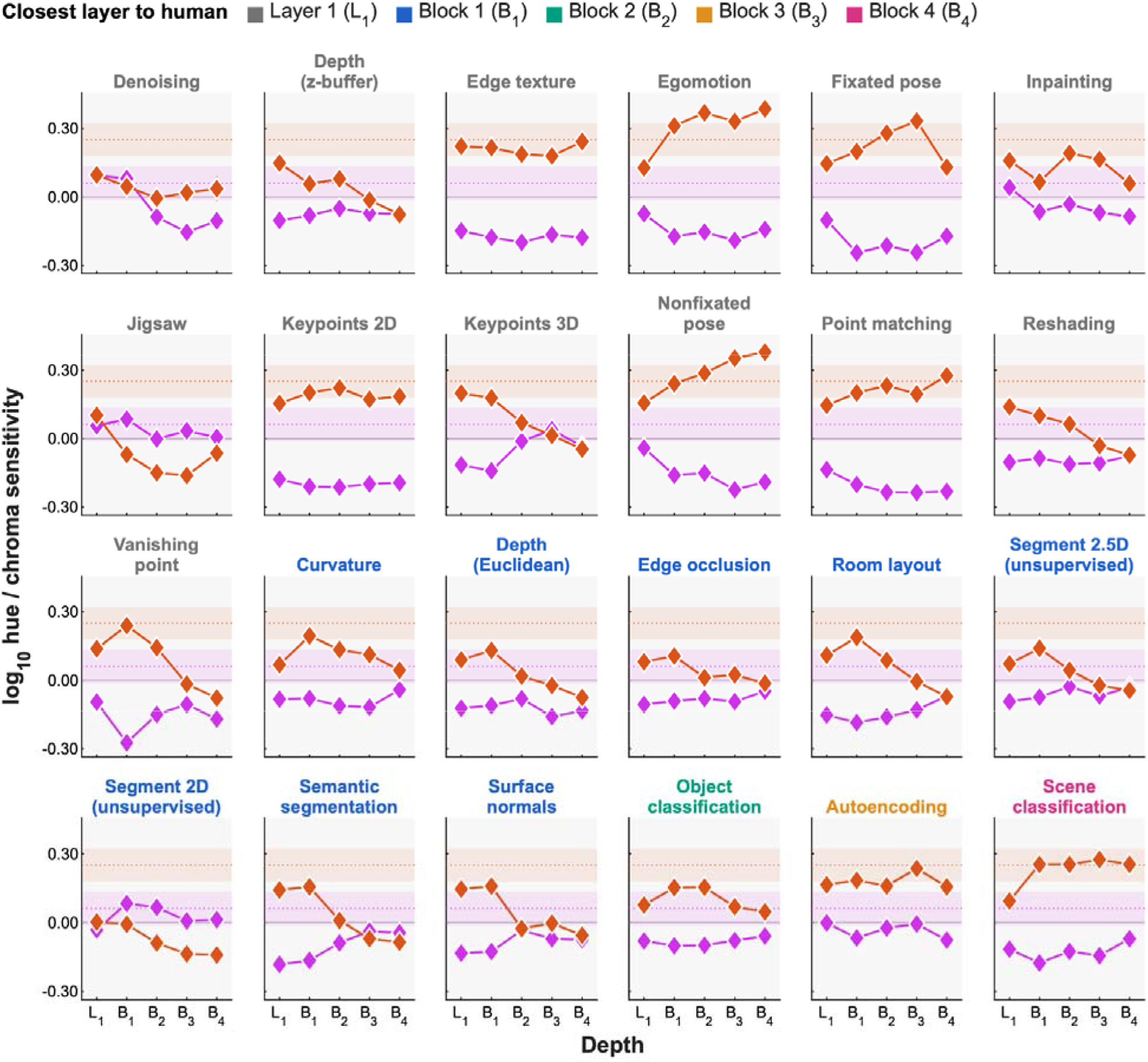
Layer-wise hue-to-chroma sensitivity ratios across Taskonomy-trained deep neural networks. Each panel shows one task; orange and magenta lines plot the log-transformed hue-to- chroma sensitivity ratio across processing depth for the orange and purple reference chromaticities, respectively. Positive values indicate higher sensitivity to hue than to chroma; negative values indicate the reverse. Shaded bands indicate the human mean ±1 SD (orange and purple), and dotted lines indicate the human means. Panel titles are colored by the best-aligned depth, matching the marker color in Figure S8.

### Layer-wise analysis of hue–chroma sensitivity in deep neural networks

We asked how the hue–chroma asymmetry evolved across processing depth in each network presented in the main text. For each layer, we computed the log-transformed hue-to-chroma sensitivity ratio separately for the orange and purple quadrants. As shown in *Figure S10*, we compared these layer-wise ratios with the human average and ±1 SD range. Each column shows networks trained on a different visual task and dataset.

All networks, except for the one trained on an S-axis-flipped ImageNet, generally showed stronger hue superiority for orange than for purple, following the human trend, but the depth at which this effect was most pronounced differed across tasks. For purple, layers from all networks except the S-axis-flipped model showed hue–chroma sensitivity ratios close to parity, with a slight bias toward chroma superiority. In the ImageNet object recognition networks, orange hue superiority was strongest in the early-to-intermediate layers (L1, B1–B3) and then declined toward the deeper, more task-specific layers; this decline was present in both ResNet50 and, more steeply, ResNet18. The COCO-trained models, which we trained from scratch, were more heterogeneous: the object detection network followed the same early-to-late decline, with its orange ratio falling close to or below zero in the deeper layers, whereas the pose estimation network showed a different profile, with an orange hue superiority that is flat or might be slightly stronger in the deeper layers. The S-axis-flipped ImageNet model showed a reversal of the orange–purple relationship.

In contrast, both Places365-trained networks maintained a high orange hue-to-chroma sensitivity ratio across a broad range of processing stages, including the deeper layers. This suggests that, for scene classification, chromatic information itself may remain directly useful for the task. Scene categories often contain diagnostic color regularities, such as sky, vegetation, water, indoor illumination, and ground surfaces, so these color statistics may remain relevant even in late representations optimized for scene identity.

This interpretation is consistent with previous work showing that CNN representations change systematically across depth. Early layers often encode generic features such as color blobs and oriented filters, whereas deeper layers become increasingly task-specific (Yosinski et al., 2014). Visualization studies similarly show a progression from low-level edges and colors to intermediate motifs and higher-level object parts (Zeiler & Fergus, 2014). Because receptive fields also expand with depth, intermediate layers can integrate color signals over larger spatial scales while still preserving information about object- or region-level structure (Araujo et al., 2019). This may explain why chromatic biases are often most prominent in shallow-to-middle layers: these stages combine relatively direct access to image color statistics with spatial scales that are large enough to capture object-relevant structure.

To quantify this progression over layers, we asked how this orange hue-to-chroma ratio changed across network depth. The S-flipped ImageNet model was excluded from this comparison because it was trained on intentionally altered chromatic statistics and showed the predicted opposite pattern. For each network, we compared an early-to-intermediate bin, comprising layer 1 and residual blocks 1–3, with the remaining deeper layers.

On average, the networks trained to perform object-level tasks showed numerically stronger orange hue superiority in the early-to-intermediate layers than in the deeper layers, but this difference was not statistically reliable and varied considerably across networks. For the ImageNet object recognition and COCO object detection and pose estimation models, the orange ratio in layer 1 and blocks 1–3 (mean = 0.222, standard deviation = 0.113) was numerically higher than in the later layers (mean = 0.159, standard deviation = 0.182), but the difference was small and non-significant (mean difference = 0.063, t(3) = 0.85, p = 0.46, 95% CI [−0.173, 0.299]). This heterogeneity was driven largely by the pose estimation network, which alone showed stronger orange superiority in its deeper layers, counter to the early-to-late decline seen in the object recognition and object detection networks. Thus, although several object-level networks expressed the orange hue advantage most strongly in early-to- intermediate representations, this was not a uniform property of object-level tasks.

**Figure S10.**
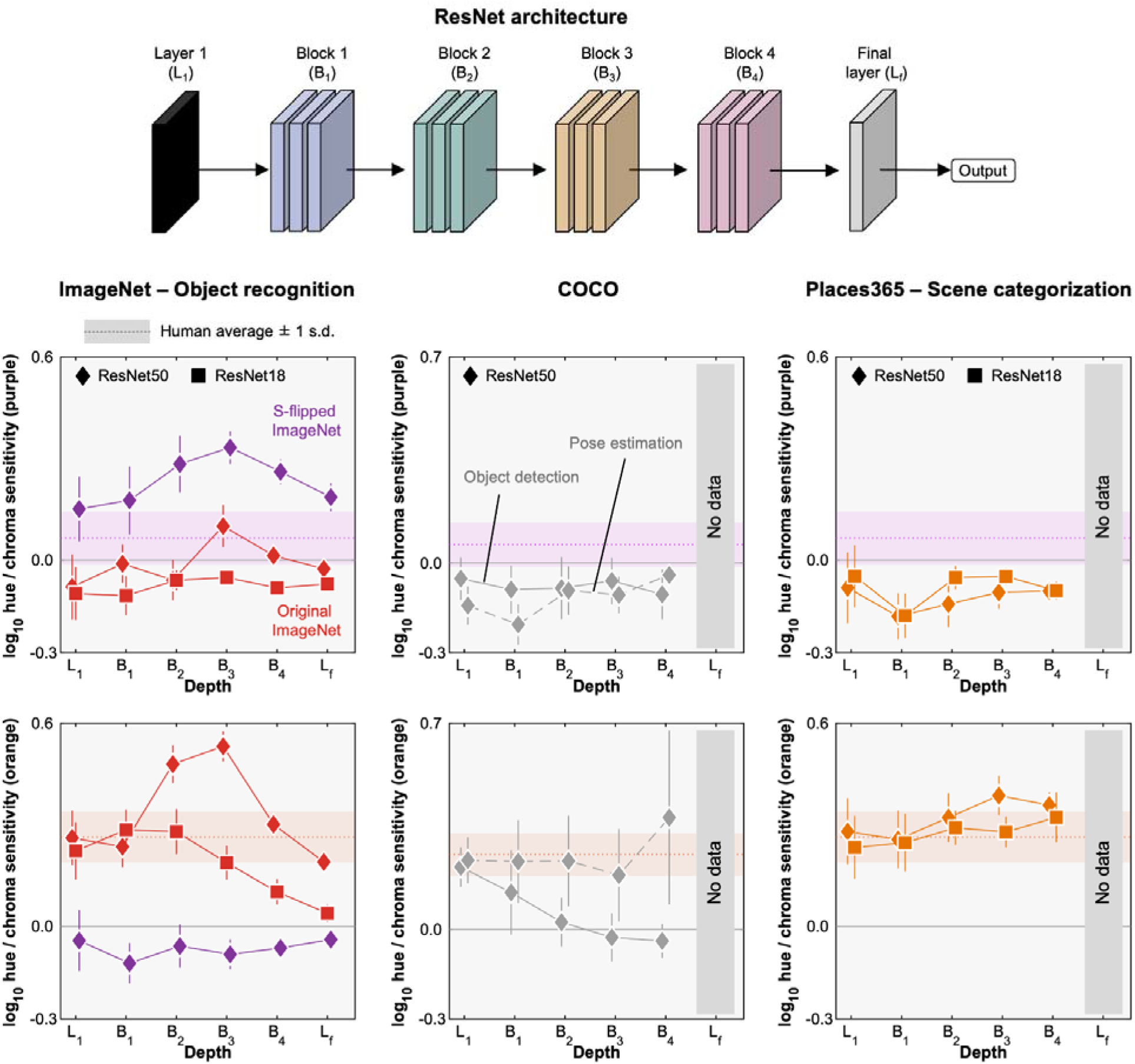
Layer-wise hue-to-chroma sensitivity ratios across deep neural networks. Log- transformed hue-to-chroma sensitivity ratios are plotted across processing depths for networks trained on different visual tasks and image datasets. Positive values indicate higher sensitivity to hue than to chroma; values near zero indicate comparable hue and chroma sensitivity. The top row shows ratios for the purple quadrant, and the bottom row shows ratios for the orange quadrant. Columns show network *trained on ImageNet for object recognition, Places365 for scene categorization, and COCO for object detection or pose estimation. Diamonds indicate ResNet50-based models, and squares indicate ResNet18-based models. Error bars show ±1 SD across training iterations. Shaded bands indicate the human mean ±1 SD for the corresponding quadrant, and dotted horizontal lines indicate the human mean*.

Scene categorization networks showed a different pattern. In the Places365-trained networks, orange hue superiority remained high across depth. If anything, the deeper layers showed a numerically larger orange hue-to-chroma ratio than the early-to-intermediate layers (layer 1 and blocks 1–3: mean = 0.273, standard deviation = 0.033; later layers: mean = 0.323, standard deviation = 0.024), although this difference did not reach significance, t(1) = −8.58, p = 0.074, 95% CI [−0.126, 0.024]. Thus, scene categorization networks preserved a strong, human-like orange hue advantage even in deeper representations.

### Detailed procedure for estimating chromatic discrimination thresholds from network representations

The procedure used to derive chromatic discrimination thresholds from network representations is summarized in *Figure S11*. We trained stage-specific linear readout classifiers to identify the odd-colored item in a four-alternative forced-choice task. The stimuli were binary masks of simple three-dimensional objects, rendered from multiple viewpoints and presented on a mid- gray background. In each trial, the four images showed the same object at different orientations. Two RGB values were drawn independently and uniformly from the RGB unit cube: one defined the color of the odd target, and the other defined the color shared by the three distractors. This sampling scheme ensured that the classifier could not rely on fixed associations between color and target position or category.

For each trial, the four images were fed into the network under evaluation. We extracted activations from multiple processing stages. When the activations retained spatial dimensions, they were averaged over space to obtain one scalar response per feature channel, yielding a compact feature vector for each image. For final vector-valued representations, such as fully connected outputs, the feature vector was used without additional pooling.

A separate linear classifier was attached to each selected network stage. On each trial, the classifier received information from the four items. This information included each item’s feature vector relative to the mean feature vector across the four items, as well as the magnitude of that deviation. Using these inputs, the classifier produced a decision score for each of the four item locations. The classifiers were trained to assign the highest score to the odd-colored target. Stage-wise losses were summed during optimization, allowing all readouts to be trained simultaneously on the same set of trials. Optimization used cross-entropy loss and the Adam algorithm.

Training was performed using 10,112 four-alternative forced-choice trials per epoch, with a batch size of 128, for 20 epochs. For each iteration, the 1,188 object shapes were partitioned into non-overlapping training and validation sets, with 90% used for training and 10% held out for testing. This procedure was repeated for 10 independent random seeds for each network architecture. The learning rate was set to 10^-3^ and weight decay to 10^-4^.

Following training, chromatic thresholds were estimated from classifier performance on the held- out validation shapes. We used the QUEST adaptive staircase procedure (Watson & Pelli, 1983), matching the approach used in the human psychophysical experiment. Thresholds were measured around two reference chromaticities, orange and purple, located on the isoluminant plane at the same radius as in the psychophysical measurements. For each reference, discrimination was assessed along both the radial chroma direction and the tangential hue direction. The chromatic coordinate system and stimulus procedure were matched to those used for the human observers.

**Figure S11.**
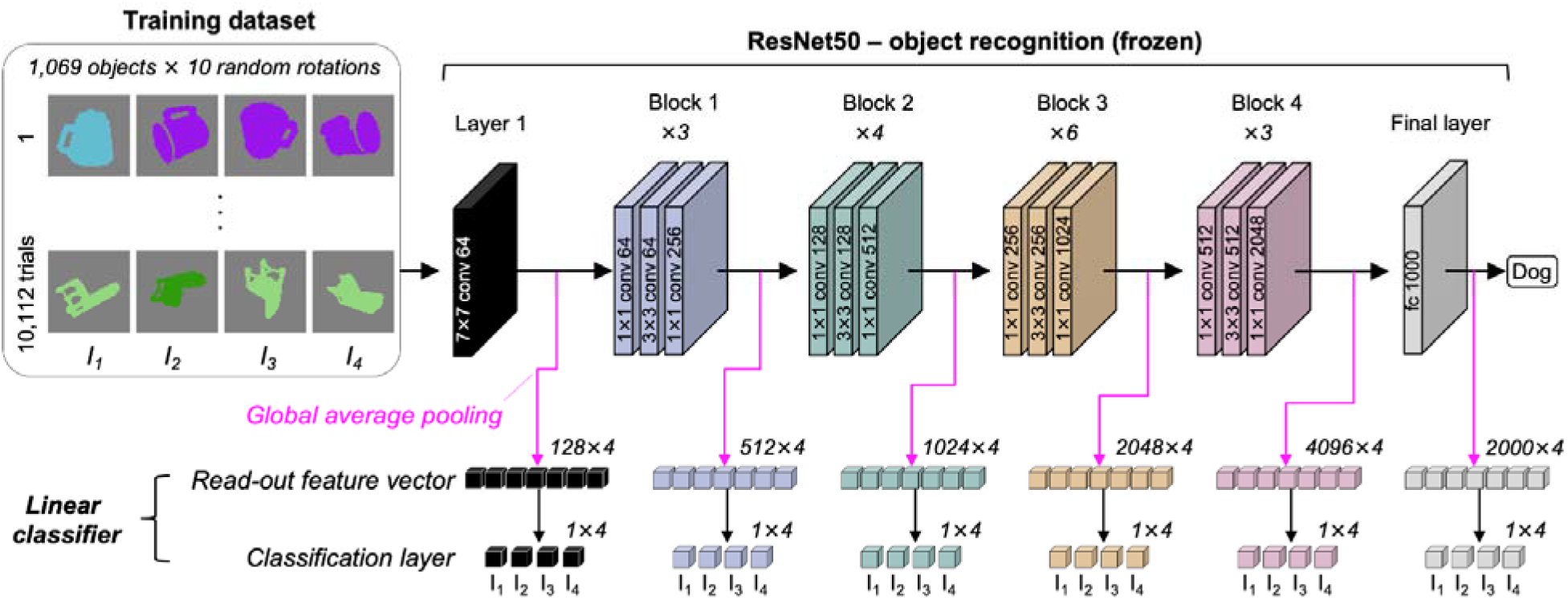
Schematic of the procedure used to estimate chromatic thresholds at different depths of a pre-trained ResNet50 (trained on ImageNet for object recognition). Responses were extracted from intermediate layers when colored 2-D shapes at different orientations were presented to the network, and these signals were used to train a linear classifier on a four-alternative forced-choice (4AFC) discrimination task. Importantly, the network was kept frozen so that its internal chromatic representations were not altered. The RGB colors of the 2-D shapes were randomly sampled to avoid introducing bias.

